# Visual–Acoustic Thigmotaxis in Zebrafish Larvae: A High throughput NAM for Neurotoxicity Assessment

**DOI:** 10.64898/2026.02.03.703464

**Authors:** Monica Torres-Ruiz, Maria Muñoz Palencia, Antonio De la Vieja, Ana I. Cañas-Portilla

**Affiliations:** Environmental Toxicology Unit, Centro Nacional de Sanidad Ambiental (CNSA), Instituto de Salud Carlos III (ISCIII), Ctra. Majadahonda-Pozuelo Km. 2,2., Majadahonda, Madrid 28220, Spain; Endocrine Tumor Unit, Unidad Funcional de Investigación en Enfermedades Crónicas (UFIEC), Instituto de Salud Carlos III (ISCIII), Ctra. Majadahonda-Pozuelo Km. 2,2., Majadahonda, Madrid 28220, Spain

**Keywords:** fish, larva, anxiety, thigmotaxis, NAM, behavior, neurotoxicity

## Abstract

The nervous system is highly vulnerable to chemical disruption, yet current regulatory guidelines do not include behavioral endpoints that capture changes in stress-related responses. Zebrafish larvae, up to 5 days old, have emerged as a promising model to bridge this gap, offering genetic and neurochemical similarity to humans together with high throughput potential. In this work, we have developed and evaluated a larval thigmotaxis assay as a new approach methodology (NAM) to detect behavioral alterations caused by neuroactive substances. Thigmotaxis, or edge-preference behavior, was studied in zebrafish larvae exposed to a range of model compounds and challenged with both visual (light/dark) and acoustic (sound/silence) stimuli. We compared 24 round well plates, commonly used in behavioral assays, with 96 square well plates to increase throughput. The two formats showed equivalent results, supporting the use of the higher-capacity system. Classical controls confirmed assay performance with caffeine increasing thigmotaxis, while diazepam decreased it. Additional neuroactive substances with diverse modes of action (chlorpyrifos, nicotine, dexamethasone, ethylenethiourea) produced stimulus-dependent responses, whereas negative controls (saccharin, amoxicillin) had little or no effect. Benchmark dose modeling showed that thigmotaxis was generally more sensitive than traditional locomotor activity endpoints. Overall, this multiplexed visual–acoustic thigmotaxis assay proved reproducible, scalable, and sensitive. In neurotoxicity testing this method could be used both as a stand-alone assay or as part of a broader behavioral NAM battery to assess potential effects on the vertebrate nervous system. This method provides a practical and ethical tool to improve chemical safety assessment both in ecotoxicology and human toxicology.

**GRAPHICAL ABSTRACT:** 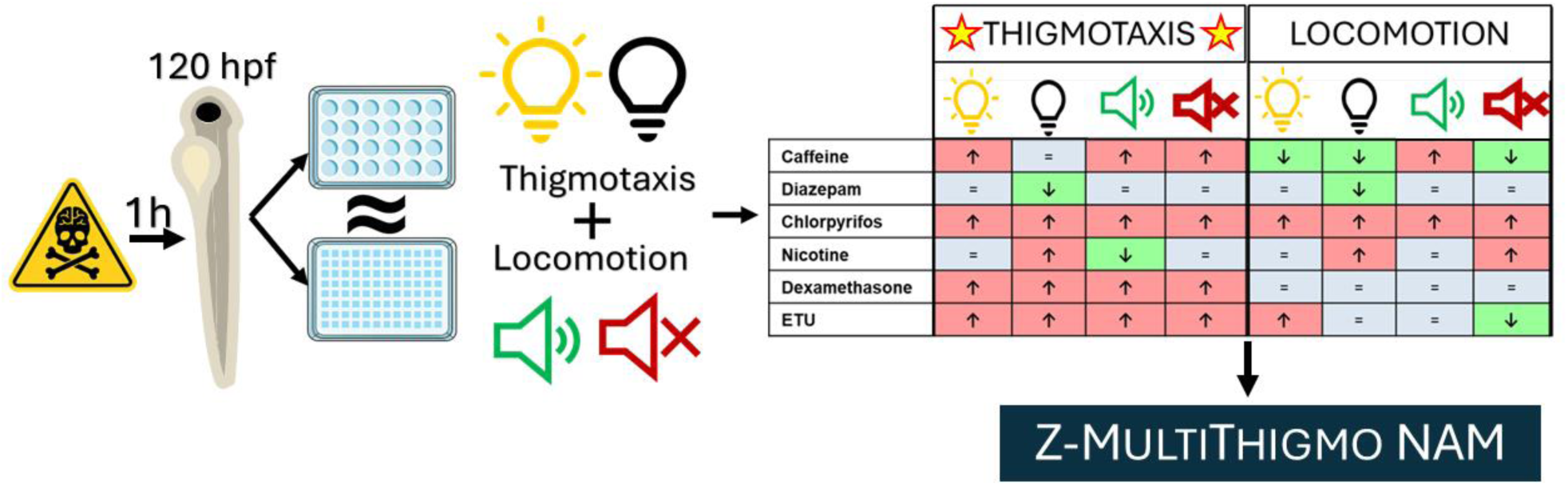

## 1. Introduction

The nervous system can be particularly susceptible to alterations by chemical substances. In the last decades the neurotoxicity (NT) and developmental neurotoxicity (DNT) potential of a great number of substances used as medical drugs, pesticides, or additives to commercial products has been demonstrated (Deepika et al., 2020; Diav-Citrin, 2011; Naranjo et al., 1981; Richardson et al., 2019). However, thousands of chemicals in the market have yet to be evaluated and some estimates predict that up to 30% may have neurotoxic potential (Legradi et al., 2018). DNT arising from insult to the developing nervous system, can have long lasting consequences and some studies have linked exposure to low concentrations of chemicals to childhood diseases such as autism spectrum or attention deficit disorders, intellectual disabilities, or learning difficulties (Grandjean and Landrigan, 2014). On the other hand, child, adolescent, and adult NT can also occur after central nervous system development due to chronic or acute chemical exposure and this in turn has been suggested to be implicated in neurodegenerative diseases such as Alzheimer’s or Parkinson’s (Chin-Chan et al., 2015). Moreover, mental disorders common in adolescents and adults, such as depression and anxiety, could also have a trigger in environmentally released neuroactive substances (Dickerson et al., 2020; Zundel et al., 2022).

The prevalence of anxiety has dramatically increased in the last decades and it is estimated at around 4% of worldwide population (Javaid et al., 2023). Anxiety and depression are a risk factor for suicide (Ned H. Kalin, 2021) and are associated with a loss of global economic productivity amounting to approximately 1 trillion US dollars per year (The Lancet Global, 2020). Despite the importance of anxiety disorders and their potential relationship to environmental pollutants, current OECD guidelines (424 and 426) to evaluate chemical neurotoxicity in a regulatory context focus on behavioral endpoints such as activity, learning, and memory but do not include anxiety-related endpoints (OECD, 1997; OECD, 2007). Moreover, rodents are the model of choice in these guidelines and therefore these studies are time-consuming, costly, and present ethical issues regarding animal welfare (Smirnova et al., 2014). These limitations of current test guidelines are stimulating the development of new approach methodologies (NAMs) to evaluate neurotoxicity. A battery of *in vitro*, cellular based, assays has been recently proposed to evaluate DNT by (Blum et al., 2023) and is in the process of being refined for regulatory use (OECD, 2023; Tal et al., 2024). Despite the ample suite of assays that compose this battery and the multiple molecular and cellular key events and modes of action that it covers, the battery has limitations. In particular, the myriads of cellular and organ interactions possibly involved in the development and maintenance of an intact nervous system and that result in complex behaviors, cannot be currently covered by solely relying on *in vitro* methods.

In this context, the zebrafish (*Danio rerio*) embryo/larvae, in addition to its wide role as an ecotoxicological model, has emerged as a powerful whole-organism translational model of human relevant hazard assessment because of its high genetic homology with humans. The zebrafish shares 71.4% of total genes, 84% of human disease genes, and 86% of human drug target genes (Gunnarsson et al., 2008; Howe et al., 2013). Moreover, zebrafish have become an excellent vertebrate model for neurotoxicity research because they retain neuroanatomical and neurotransmitter systems very similar to those of mammals and are metabolically competent (Chu and Sadler, 2009; d’Amora and Giordani, 2018; Gupta et al., 2018; Horzmann and Freeman, 2016). Primary neurogenesis in zebrafish is completed at around 72 hours post fertilization (hpf), and at 120 hpf their nervous system is mostly mature with larvae exhibiting a large repertoire of visual and acoustic induced behaviors (McAtee and Abdelmoneim, 2024; Muto and Kawakami, 2013). In addition, zebrafish larvae up to 120 hpf are considered to meet the requirements of the 3Rs principle and are not regulated as animal experiments by current legislation (Bauer et al., 2021). In this context, assays that rely on early life stage zebrafish are being proposed to be included in a second generation developmental neurotoxicity test battery and a first generation adult neurotoxicity test battery by participants of the project PARC (Partnership for the Assessment of Risks from Chemicals), cofinanced by the European Union (Tal et al., 2024). The present work, as part of this task, has the objective of setting up a high throughput zebrafish larvae thigmotaxis assay to evaluate effects of chemicals on anxiety-like behavior as a result of visual and acoustic stimuli.

Thigmotaxis, a term used to describe the movement of an organism in reaction to surfaces or objects, is a behavioral response to tactile stimuli that has been observed in evolution as early as in protozoans (Ricci, 1990). When placed in a new confined location, for instance, animals often stay toward the periphery and avoid the interior. Thigmotaxis has been extensively used as a measure of anxiety (Prut and Belzung, 2003) but it is also possible that edge preference could be caused by alternative mechanisms such as sensory deficits, motor impairment, or hyperexcitability caused by chemical exposure (Bellot et al., 2024; Gerlai, 2016). Nevertheless, regardless of the specific mode of action of thigmotaxis, it is clear that this conserved behavior is a good candidate to evaluate effects of neuroactive substances. Proof of this is the fact that this wall-hugging innate response has been used extensively in rodent studies as a measure of behavioral change in relation to drug or chemical exposure (Estrela et al., 2021; Karl et al., 2003; Prut and Belzung, 2003). Moreover, it has also been used in humans as a measure of anxiety (Gromer et al., 2021). In zebrafish larvae, thigmotaxis (or open field tests) studies have been used mainly to characterize neurological disease models (Christensen et al., 2020), as an aid to find new anxiolytic substances (Muniandy, 2018), or to test toxicant effects (Gomes et al., 2024; Richendrfer et al., 2012b). However, when studying effects on zebrafish larvae, thigmotaxis has been experimented with a general lack uniformity, with a wide variety of test conditions and endpoints assessed. Moreover, studies rarely use adequate positive/negative controls (Liu et al., 2016; Martins Fernandes Pereira et al., 2022; Yang et al., 2017). Most works also have a relatively low throughput, using mainly 6,12, or 24 well plates, and rarely evaluate thigmotaxis responses to changing visual cues or, even less, to acoustic stimuli (Lindemann et al., 2022; Merola et al., 2021b; Richendrfer et al., 2012a; Schnörr et al., 2012).

Therefore, the **objective** of the present work was to develop and assess the performance of a high throughput zebrafish larvae (120 hpf) thigmotaxis assay in order to be used in drug and toxicological assessments. Given the versatile nature of the model, this assay could be used both in the field of ecotoxicology and also in determining effects on the human nervous system. We have compared test performance in 24 and 96 well plate formats and we provide detailed information on normal and solvent (DMSO) control thigmotaxis responses during periods of visual and acoustic stimulation. In addition, we have tested neurotoxic model substances to establish method readiness and to study the added value of a thigmotaxis assay as it compares to the more common test of light/dark transition as a measure of neuroactivity and/or anxiety (Dach et al., 2018; Irons et al., 2010). This assay will be later validated in a regulatory context to be used as part of a neurotoxicity assessment battery.

## 2. Material and Methods

### 2.1. Zebrafish husbandry

Adult wild-type zebrafish (AB strain) were housed in 30 L glass recirculating aquariums with a temperature of 25–26°C, a pH of 7–7.5, oxygen levels of 6.5–7 mg/L, and conductivity ranging from 500 to 600 μS/cm. The fish water was prepared using deionized water from a MilliQ system (Merck RiOs™ Essential 24) and a mixture of salts: CaCl_2_·2H_2_O (294 mg/L), MgSO_4_·7H_2_O (123 mg/L), NaHCO_3_ (64.7 mg/L), and KCl (5.8 mg/L). The fish were kept on a 14:10 h light/dark cycle and were fed twice daily with frozen brine shrimp (Artemia salina, Ocean Nutrition™, Belgium), shell-free brine shrimp eggs (Ocean Nutrition™), or Tropica Basic fish flakes (Dajana®, Czech Republic).

### 2.2. Embryo collection and culture

Embryos were obtained by random mating of adult zebrafish. The afternoon before, 2 females and 3-4 males were placed in a sloping breeding tank with a perforated bottom (Tecniplast), separated by a removable divider. In the morning, as soon as lights were turned on, the divider was removed, and animals were allowed to mate for 1 hour undisturbed. Embryos were then collected and washed gently 4 times with egg water (same as fish water, see above) to remove debris. Between 2-3 hpf, embryos were carefully observed under an Ivesta 3 stereoscope (Leica Microsystems) and only embryos showing proper development were chosen and placed in 100 mm petri dishes (50 embryos/plate). Embryos were incubated at 28±0.5°C and 14:10 h light/dark photoperiod (lights on at 6:00 am) in a highly controlled climate chamber for 96 h (Fitoclima 1200, Aralab). No ethics approval was required as zebrafish larvae reared in our institution do not exhibit independent feeding at the stages when the experiments were terminated (around 120 hpf) (EU Directive 2010/63/EU). Nevertheless, embryos were treated according to humane principles, always manipulated with care using wide tip plastic pipettes and euthanized using tricaine methasulfonate MS-222 (160 mg/L; adjusted to pH 7.0–7.5 with Tris Buffer pH 8) and later freezing at -20°C upon experiment ending or if signs of malformations or stress were observed during development.

### 2.3. Exposure regimes

#### 2.3.1. Plate selection

Two types of plates were tested, Falcon 24 round (Catalog number 353047) and Whatman 96 square well plates (Catalog number 7701-1651) due to availability from plate manufacturers (24/48 square well plates do not exist in the market). The 24 round plates (24R) are the most prevalent among zebrafish larvae thigmotaxis studies (Schnorr et al. 2012) but provide low throughput. In addition, we had previously observed a higher degree of variability when using round plates vs square plates for zebrafish larvae behavioral analyses. To test the possibility of increasing throughput we chose a 96 well format, and the square configuration was chosen due to greater surface area than traditional round wells and because we hypothesized this format would decrease behavioral variations.

#### 2.3.2. Acclimation and Toxicant Exposure

The assay has been developed for an acute neurotoxicity mode, meaning that larvae were exposed for a short period of time (1h) at 120 hpf when most organs are already developed.

Trying to reduce possible disturbance sources on the exposure day (120 hpf), we randomly transferred larvae the day before, at 96 hpf, from the 100 mm petri dishes to 24 round (24R) and 96 square (96S) wells plates, placing one larva per well and setting 12 larvae per condition (Fig. 1). In the case of the 24R well plates each well was previously filled with 1 mL of fish culture water and 400 µL were placed in each well of the 96S well plates. Plates were sealed with parafilm to avoid water evaporation and placed in the climate chamber.

**Fig 1.**
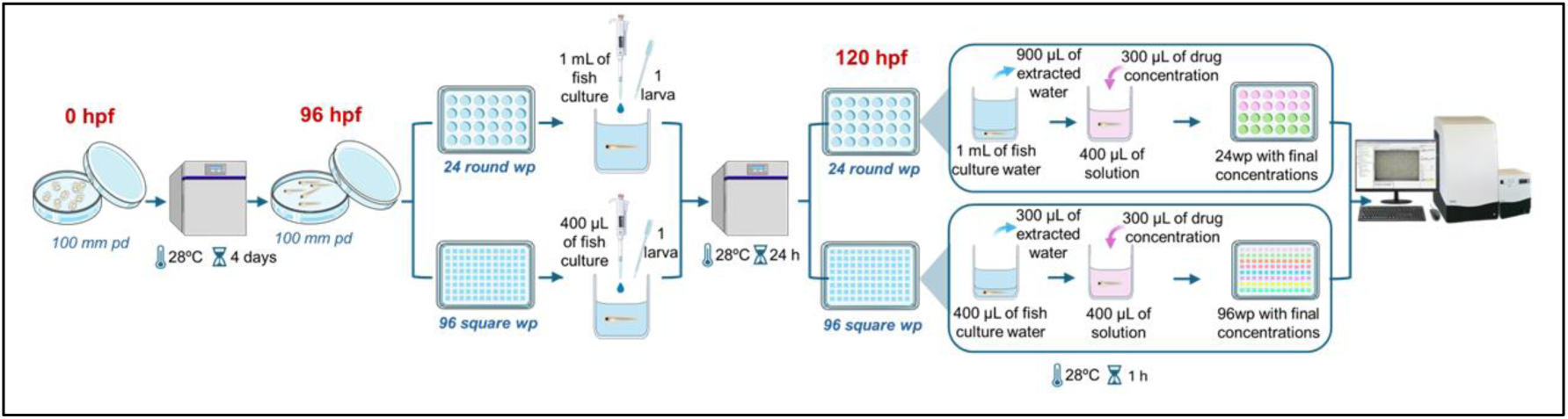
Schematic representation of culture, acclimation and exposure of larvae to chemical solutions.

After 24 hours of acclimation in the plates, at 120 hpf, we partially replaced fish water of the plate’s wells with experimental solutions (900 µL from 24R well plates and 300 µL from 96S well plates) and refilled them with different drug concentrations (Fig. 1). Final volume in each well for both plate formats was 400 µL. When appropriate, DMSO was used as a vehicle to dissolve some of the substances in a concentration no greater than 0.4 % as we observed that concentrations up to 1% had no effect on thigmotaxis after acute 1 h exposures at 120 hpf (Fig. A.1). Calculations were made so that final treatment concentrations were equal for both plate formats. Plates with larvae were then incubated for one hour in the climate chamber at 28 °C under light conditions and were then immediately evaluated for their behavior.

In order to test the performance of 24R vs 96S well plates we chose two substances that have been used widely to validate anxiety assays, caffeine as an anxiogenic and diazepam as an anxiolytic (Dos Santos et al., 2024; Richendrfer et al., 2012a; Stewart et al., 2011). These two substances were dissolved in fish water and DMSO respectively, and a range of concentrations were assessed (Table 1). In addition, other substances were tested to test assay sensibility to neurotoxicants (Chlorpyriphos, Nicotine, Dexamethasone, and Ethylenethiourea) and potential negatives (Amoxicillin and Saccharin). All compounds were obtained from Sigma–Aldrich (Zwijndrecht, The Netherlands).

**Table 1.** Substances and concentrations used in the present study, CAS number in parenthesis. All were dissolved in fish water plus a ≤ 0.4% percentage of DMSO in indicated cases. Concentration ranges used are indicated in mg/L, and maximal concentration also in µM. Last column indicates the number of replicates for each substance.

| Substance | Vehicle | Concentrations | Replicates |
| --- | --- | --- | --- |
| Caffeine (58-08-2) | Fish water | 1.0 - 100 mg/L (515 $\mu\text{M}$) | 24 |
| Diazepam (439-14-5) | DMSO | 0.001 - 10 mg/L (35 $\mu\text{M}$) | 24 |
| Chlorpyrifos (2921-88-2) | DMSO | 0.3 - 100 mg/L (285 $\mu\text{M}$) | 24 |
| Nicotine (54-11-5) | DMSO | 0.3 - 100 mg/L (617 $\mu\text{M}$) | 24 |
| Dexamethasone (50-02-2) | DMSO | 0.3 - 100 mg/L (255 $\mu\text{M}$) | 24 |
| Ethylenethiourea (96-45-7) | DMSO | 0.3 - 100 mg/L (979 $\mu\text{M}$) | 24 |
| Amoxicillin (26787-78-0) | DMSO | 0.01 - 150 mg/L (819 $\mu\text{M}$) | 24 |
| Saccharin (81-07-2) | DMSO | 0.01 - 150 mg/L (410 $\mu\text{M}$) | 12 |

#### 2.3.3. Behavioral tests

Larvae (120 hpf) were put into 24R and 96S well plates, one larva per well. For the 96 well format, the total arena area (well size) was 36 mm^2^ (6 x 6 mm) and the inner (center) and outer (edge) zones measured 16 mm^2^ (4 x 4 mm) and 20 mm^2^ respectively (Fig. 2A). The inner zone area was chosen for best performance after testing different measurements of the inner zone ranging from 3 mm^2^ to 5 mm^2^. For the 24-well format, the total arena area was 177 mm² (diameter 15 mm), and the inner and outer zones measured 80 mm² (diameter 10 mm) and 97 mm², respectively. For both plate formats the inner zone was 45 % of the total well area.

**Fig 2.**
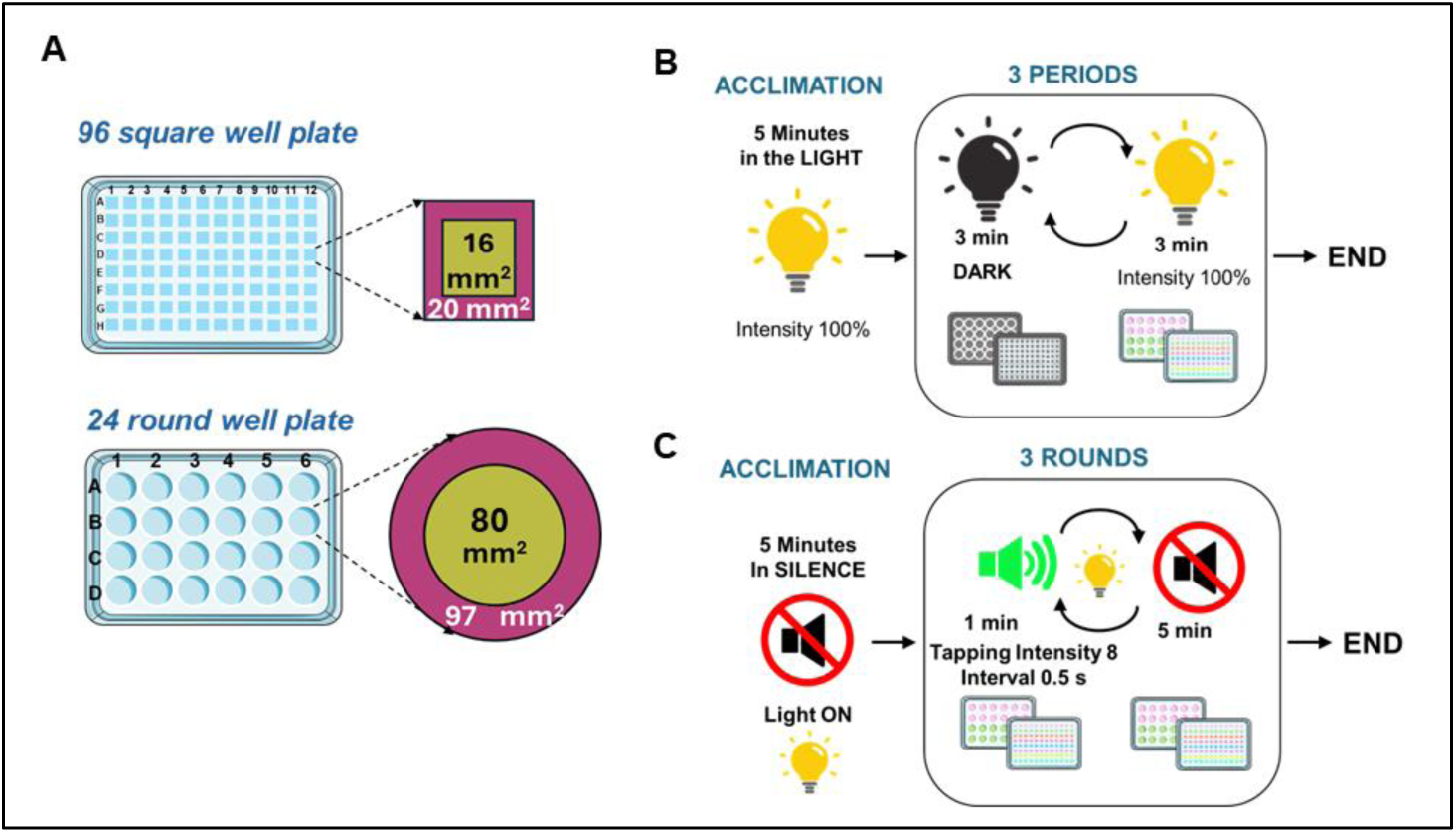
Plate geometry, sizes and stimuli regimes. (A) Size of 96 square well plates (96S) and 24 round well plates (24R) wells showing center (green) and edge (purple) zones. Schematic representations of visual (B) and acoustic (C) stimuli parameters

Behavioral assessments were carried out using the Danio Vision platform (Noldus, Wageningen, the Netherlands), and a Danio Vision Observation Chamber (DVOC) equipped with near-infrared illumination and a temperature control unit to maintain the system at 28 ◦C during all trials. Head point detection in EthoVision XT v. 16 (Noldus, The Neatherlands) was used to define the animal’s position. This approach effectively excludes the tail region from tracking, providing a more accurate estimate of the larva’s proximity to the well edge in both 24R and 96S plate formats. Recordings were made at a frame rate of 30 fps. All assays were performed at least 4 hours after daytime started for the larvae (6 am) and had the same routines for both plate formats.

For measuring anxiety responses, we exposed the larvae to two stressful conditions, first to visual stimuli, consisting of short dark/light transitions and later to acoustic stimuli consisting of a battery of tapping noises followed by silent periods (Fig. 2B,C). The visual stimuli assay started with a period of acclimation lasting 5 min in the light. Without delay, it was followed by a dark/light routine consisting of three periods of lights-off (duration 3 min) intercalated by three periods of lights-on (duration 3 min; intensity 100%). The acoustic stimuli assay was performed in the light. It started with a 5 min acclimation period in silence, followed by 3 rounds of tapping sounds (intensity 8: approximately 80-90 dB re 20 µPa in air above plate) in 0.5 second intervals with a duration of 1 minute. In between each tapping round (“tapp”), there was a 5-minute silent period (“no tapp”). After behavioral tests were finished, larvae were euthanized using rapid cooling (Wallace et al., 2018).

The Ethovision XT v. 16 software was used to analyze videos, and the following parameters were recovered: total distance moved in all the well and time spent at the edge of well for light/dark and tapp/no tapp periods.

### 2.4. Data Analysis

Thigmotaxis was assessed by % time spent by larvae in the edge of the plate relative to controls according to the formula

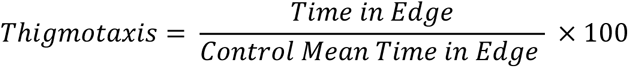

This was calculated for periods of light and dark for the visual response experiment and during tapping (tapp) and silent (no tapp) periods for the acoustic response experiment. In addition, total distance moved by larvae during light, dark, tapp and no tapp periods were assessed for all experiments. Larvae presenting with little (< 10 % distance traveled with respect to control), or no locomotion were not used for analysis because swimming movement is necessary for an unbiased measurement of zone preference (Bouwknecht and Paylor, 2008). In addition, in order to decrease natural high standard deviations common to the zebrafish model of behavior (Fitzgerald et al., 2019), the data curation included removing of the highest value and lowest value of controls and each of the compounds concentrations. Means were not significantly altered by this data curation process.

For statistical analysis, differences in thigmotaxis between plate formats and between controls and treated larvae were assessed using Two-way ANOVA with Sidak’s multiple comparisons test using GraphPad Prism 10.2.3 (*p* ≤ 0.05).

Benchmark dose (BMD), with upper and lower limits, was calculated for plate format comparison and for endpoint comparison (thigmotaxis vs total distance moved) using the Benchmark Dose Online Tool (BMDS found at https://bmdsonline.epa.gov/; Environmental Protection Agency, USA) for cases in which significant differences were found between control and exposure doses. The benchmark response (BMR) was defined as a change of 1 standard deviation from the control mean. A one-sided 95% confidence level (tail probability = 0.05) was used to estimate the lower confidence limit of the BMD. In cases where the dose response was not monotonic data for some concentrations (generally the highest) were eliminated from analysis. In all cases, the Hill model was the most appropriate for the reported data. In a handful of cases the BMDS tool was unable to fit a model to the data, and the BMD was not calculated.

The bioequivalence of 24 round to 96 square well plates was evaluated in Microsoft Excel using a Two One-Sided Test (TOST) to determine if it is possible increase throughput while preserving the interpretability of thigmotaxis readings (Schuirmann, 1987; Walker and Nowacki, 2011). The TOST allows a positive conclusion of practical equivalence when the 90% confidence interval (CI) of the format difference lies within an *a priori* margin of indifference. This strategy is standard in bioequivalence and method transfer and is increasingly used for assay interrelationships (Aabeyir et al., 2020). Moreover, it is preferred over the difference test to validate whether two assay formats produce substantially the same result within a prespecified tolerance and equivalence testing. In the present work, we compared readings of control larvae from 96 wells (96S) with those from 24 wells (24R) for each assay stimulus type and condition using the TOST test with α = 0.05 and an equivalence margin of Δ = ±10 percentage points (pp). We calculated the difference in means (96S − 24R) with Welch standard errors and Satterthwaite degrees of freedom and tested two one-sided hypotheses: the difference is greater than −Δ and less than +Δ, respectively. Equivalence is concluded when both tests are significant. That is, the 90% CI lies entirely within the range [−10, +10]) (Aabeyir et al., 2020; Schuirmann, 1987; Walker and Nowacki, 2011). For concentration-dependent experiments with reference substances diazepam and caffeine, where visual (light/dark) and acoustic (sound/silence) stimuli were assessed, differences by concentration were pooled within each trial and condition using a fixed-effects inverse variance meta-analysis. Satterthwaite’s minimum degree of freedom across concentrations was used to obtain conservative intervals. Decision criteria were classified as "Equivalent" (TOST *p*_max < 0.05), "Different" (Welch two-tailed *p* < 0.05), or "Inconclusive" otherwise.

### 2.6. Ethical considerations

Experiments were conducted using zebrafish embryos up to 120 hpf, so institutional approval for animal care was not required. Nevertheless, all experimental procedures were performed in accordance with international accepted standards for humane use of model organisms, and every effort was made to minimize the number of embryos used and to ensure optimal husbandry conditions.

## 3. Results

### 3.1. Plate format comparison under visual and acoustic stimulation

The initial objective was to determine whether the assay could be scaled from a 24-well round plate to a 96-well square plate in order to increase throughput. To carry out the assay, control larvae were maintained in embryo water without exposure to any substance. Control larvae behaved very similarly for both plate formats and both types of stimuli, with greater % time and distance in the edge of the wells, indicating, as expected, that they exhibit natural thigmotaxis (Fig. 3). Percent distance/zone and total distance (cm) were slightly higher in 24 well plates (only in light periods; Fig. 3B, C) as expected from more surface area but there were no significant differences in time spent in each zone with respect to plate format for both types of stimuli (Fig. 3A, D).

**Fig 3.**
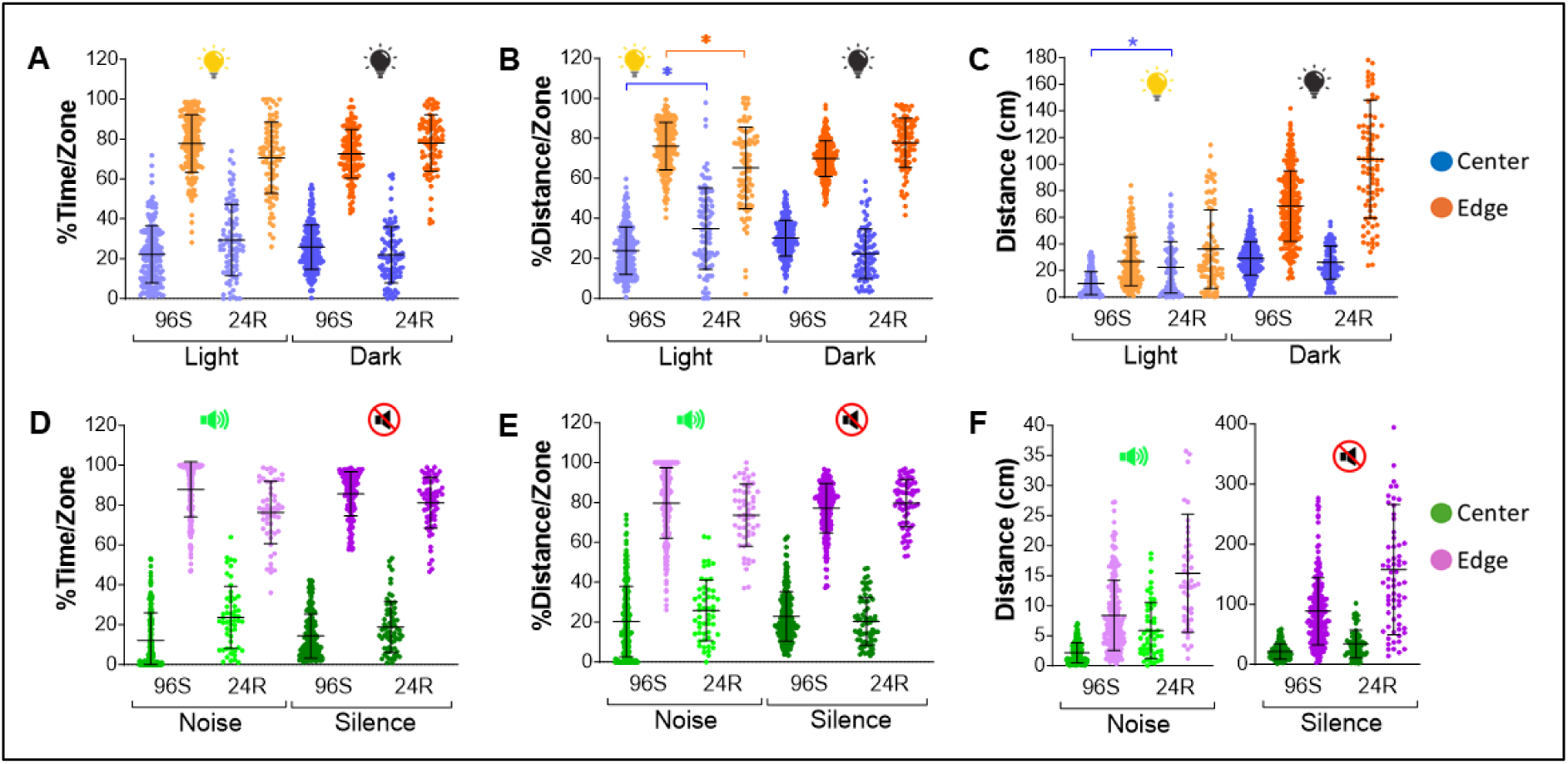
Results of control larvae for both visual and acoustic stimuli and both types of plate formats (24R and 96S). Panels A-C depict natural thigmotaxis results in response to visual stimuli, with percentage of time in each zone (A), percent distance in each zone (B) and absolute distance traveled in each zone (C) by larvae for the center of the plates in blue and the edge in orange. Panels D-F depict natural thigmotaxis results in response to acoustic stimuli, with percentage of time in each zone (D), percent distance in each zone (E) and absolute distance traveled in each zone (F) by larvae for the center of the plates in green and the edge in purple. 96S: 96 square well plates; 24R: 24 round well plates. Asterisks denote significant difference of 24R vs 96S plates (* p<0.05). Data are presented as Means +/- SD.

Having established baseline thigmotaxis performance in untreated controls, we next assessed the assay’s sensitivity by exposing larvae to reference anxiogenic caffeine and anxiolytic diazepam across both plate formats and stimulus modalities. Both caffeine and diazepam behaved similarly for both plate formats and both stimuli, with generally no significant differences in responses between plates (Fig. 4, Table 2). Larvae exposed to caffeine showed an increased thigmotaxis (20-25% at highest concentration) during light periods (Fig. 4A) but not during dark periods, except at the highest concentration for the 96S plates only (Fig. 4B). Caffeine also induced increased time at the edge during periods of tapping noises with very similar behavior in both plate formats (Fig. 4C). During silent periods, larvae in the 96S plate format exhibited a gradual dose-response curve whereas larvae in the 24R plate were more variable (Fig. 4D). Despite this, BMDs were similar for both (Table 2).

**Fig 4.**
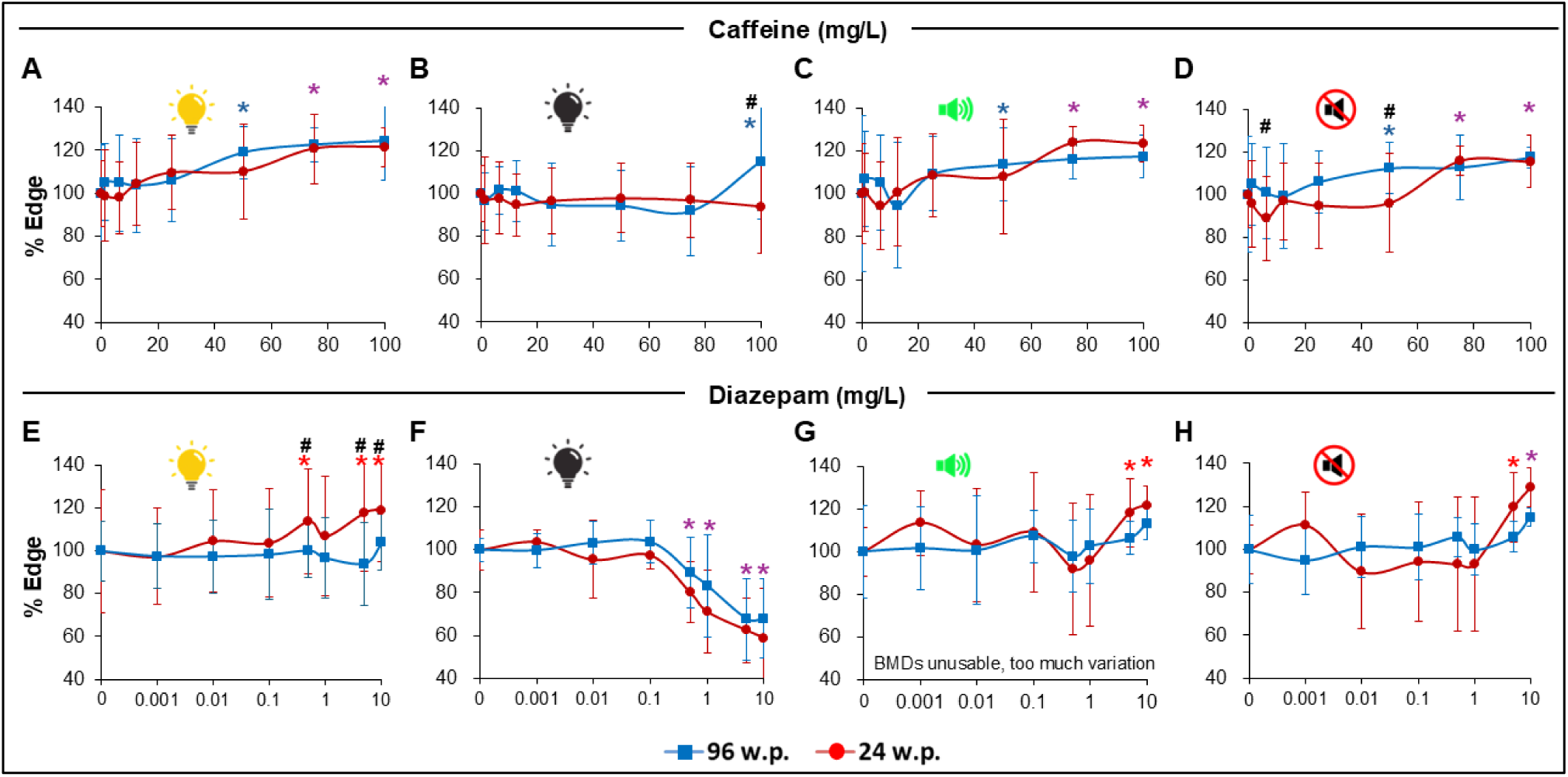
Results of larvae exposed to model anxiogenic caffeine and anxiolytic diazepam substances for both types of stimuli (visual and acoustic) and both types of plate formats (24R and 96S). Blue lines correspond to a 96 square well plates and red lines to 24 round well plates. Panels A-D depict thigmotaxis results in response to Caffeine exposure, with percentage of time in each zone during light periods (A), dark periods (B), tapping periods (C) and silent periods (D). Panels E-H depict thigmotaxis results in response to Diazepam exposure, with percentage of time in each zone during light periods (E), dark periods (F), tapping periods (G) and silent periods (H). Purple asterisk above point denotes significant differences of substance concentration vs control for both plate formats. Blue asterisk above point denotes significant difference of substance concentration vs control for 96 square well plate formats. Red asterisk above points denotes significant differences of substance concentration vs. control for 24 round well plate format. Hash symbols denote significant differences between plates. Data are presented as Means +/- SD.

**Table 2.** Benchmark dose (BMD) with lower (BMDL) and upper (BMDU) limits for larvae exposed to Caffeine and Diazepam for light, dark, tapping (Tapp), and quiet (No Tapp) periods and two types of plate formats, 24 round well plates (24R) and 96 square well plates (96S) calculated using the EPA BMD Tool (Hill model). When the BMDS tool was unable to fit a model to the data the BMD was not calculated (Model unusable as BMDS tool was unable to fit a model to the data). MOA: Primary pharmacological class or mode of action (MOA)

| Substance | MOA | Interval | Plate Format | BMDL | BMD | BMDU |
| --- | --- | --- | --- | --- | --- | --- |
| Caffeine | non-selective adenosine receptor antagonist | Light | 24R | 48.4 | 78.7 | 100.2 |
|  |  |  | 96S | 43.2 | 57.2 | 58.4 |
|  |  | Dark | 24R | Non-significant. Analysis not done |  |  |
|  |  |  | 96S | Non-significant. Analysis not done |  |  |
|  |  | Tapp | 24R | 56.3 | 61.7 | 64.6 |
|  |  |  | 96S | 62.0 | 82.7 | 84.6 |
|  |  | No Tapp | 24R | 61.8 | 67.8 | 75.5 |
|  |  |  | 96S | 25.0 | 85.8 | 89.8 |
| Diazepam | benzodiazepine GABA<sub>A</sub> receptor positive allosteric modulator | Light | 24R | 10.0 | 15.6 | 37.4 |
|  |  |  | 96S | Non-significant. Analysis not done |  |  |
|  |  | Dark | 24R | 0.14 | 0.31 | 0.48 |
|  |  |  | 96S | 0.31 | 0.42 | 0.43 |
|  |  | Tapp | 24R | Model unusable |  |  |
|  |  |  | 96S | Model unusable |  |  |
|  |  | No Tapp | 24R | 3.2 | 4.4 | 4.5 |
|  |  |  | 96S | 5.5 | 7.5 | 7.7 |

Larvae exposed to Diazepam exhibited a very similar (30-40% at highest concentration) decrease in thigmotaxis in both plate formats but only during dark periods (Fig. 4F, Table 2). For larvae in the 24R plates, an unexpected increase in time at the edge was observed during light periods. This increase was not observed in larvae tested in the 96S plates (Fig. 4E). During acoustic stimulation, larvae exhibited increased thigmotaxis at the two highest Diazepam concentrations for both plate formats (Fig. 4G). A similar pattern was observed during silent periods (Fig. 4H, Table 2).

To ensure the equivalence between plates, a TOST analysis was performed. The pooled TOST results show that most contrasts between test conditions for 96S and 24R plates are equivalent, within a strict ±10 pp difference (Table A.1 and Fig. A.2). In controls, equivalence was observed for percentage visual time and distance in cm, as well as for percentage acoustic time and distance in cm. Only percentage acoustic distance was classified as different, in logical line with the geometric mapping scaling between formats. For reference compounds, caffeine was equivalent across all visual and acoustic conditions, while diazepam was equivalent for visual dark (decreased thigmotaxis) and acoustic tap, but different for visual light (increased thigmotaxis observed only in 24R plates) and acoustic quiet.

Plate format was also tested in other neuroactive substances such as tofisopam, ethanol and valerianic acid, and non-neuroactive as ibuprofen. In general, similar responses were observed (Fig. A.3) so the 96S plate format was deemed appropriate to continue with the study of how thigmotaxis compares to general locomotion as a measure of neuroactivity.

### 3.3. Neuroactive and negative substances visual tests

We next challenged the assay with a broader panel of neuroactive substances spanning distinct pharmacological classes and modes of action (MOA). Four neuroactive model substances were used: the pesticide organophosphate cholinesterase inhibitor Chlorpyrifos, the stimulant alkaloid acetylcholine receptor agonist Nicotine, the fluorinated glucocorticoid receptor agonist Dexamethasone, and the organosulfur rubber additive thyroid-disrupting goitrogen Ethylenethiourea. In addition, two putatively negative substances with no neuroactive MOAs in the sense of CNS receptor targets relevant to anxiety assays were also tested: the benzosulfimide artificial sweet-taste receptor agonist Saccharin and the β-lactam antibiotic and cell-wall synthesis inhibitor Amoxicillin. All substances were tested in the visual and acoustic stimuli modes (Fig.5; Table 3).

**Fig 5.**
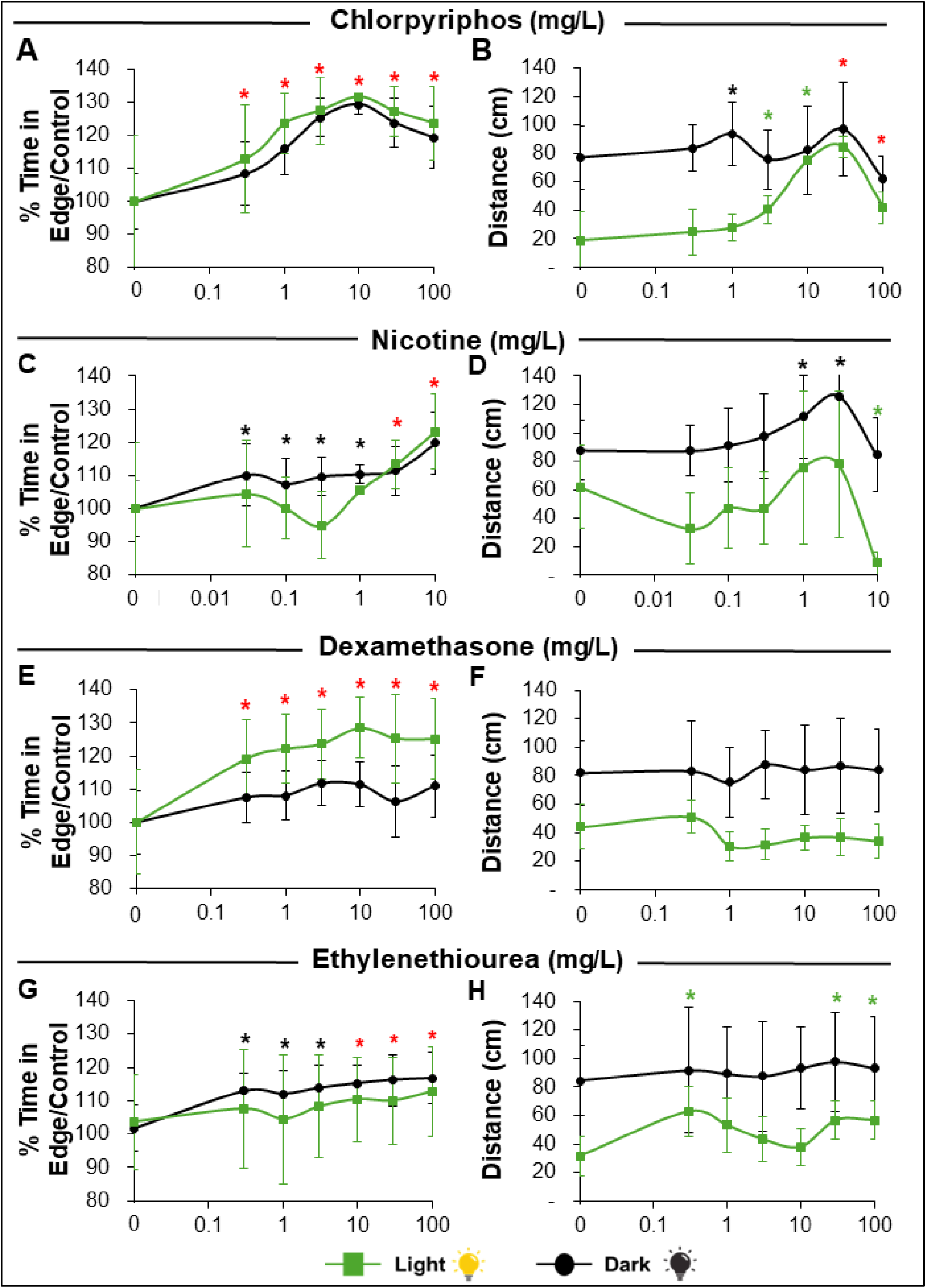
Results of Thigmotaxis and Total distance moved by larvae exposed to neurotoxic substances (chlorpyriphos, nicotine, dexamethasone and ethylenethiourea) in 96S well plates after visual stimulation. Green lines depict light periods, and black lines depict dark periods. Green asterisk above point denotes significant differences of substance concentration vs. control during light period. Black asterisk above point denotes significant differences of substance concentration vs. control during dark period. Red asterisks above points denote significant differences of substance concentration vs. control for both periods. Data are presented as Means +/- SD.

**Table 3.** Benchmark dose (BMD) with lower (BMDL) and upper (BMDU) limits for larvae exposed to different neuroactive substances for thigmotaxis and distance travelled during Light, Dark, tapping (Tapp), and quiet (No Tapp) periods calculated using the EPA BMD Tool (Hill model). When the BMDS tool was unable to fit a model to the data the BMD was not calculated (Model unusable as BMDS tool was unable to fit a model to the data). NC: not calculated, BMDS tool could not calculate an upper limit. MOA: Primary pharmacological class or mode of action (MOA)

| Substance | MOA | Endpoint | Interval | BMDL | BMD | BMDU |
| --- | --- | --- | --- | --- | --- | --- |
| Chlorpyrifos | Organophosphate cholinesterase inhibitor | Thigmotaxis | Light | 0.2 | 0.3 | 0.6 |
|  |  |  | Dark | 0.2 | 0.4 | 0.6 |
|  |  | Distance | Light | 1.7 | 3.0 | 4.8 |
|  |  |  | Dark | 22.1 | 43.0 | 143.3 |
| Nicotine | Nicotinic acetylcholine receptor agonist | Thigmotaxis | Light | 1.9 | 3.3 | 6.1 |
|  |  |  | Dark | 0.05 | 0.2 | 1.9 |
|  |  | Distance | Light | 0.5 | 0.8 | 1.2 |
|  |  |  | Dark | 0.3 | 0.8 | 1.8 |
| Dexamethasone | Glucocorticoid receptor agonist | Thigmotaxis | Light | 0.06 | 0.15 | 0.6 |
|  |  |  | Dark | 0.1 | 0.3 | 0.4 |
|  |  | Distance | Light | Non-significant. Analysis not done |  |  |
|  |  |  | Dark | Non-significant. Analysis not done |  |  |
| Ethylenethiourea | Thyroid-disrupting goitrogen | Thigmotaxis | Light | 10.3 | 10.8 | 14.8 |
|  |  |  | Dark | 0.1 | 0.2 | 0.3 |
|  |  | Distance | Light | Model unusable |  |  |
|  |  |  | Dark | 30.0 | 62.2 | 371.0 |
| Chlorpyrifos | Organophosphate cholinesterase inhibitor | Thigmotaxis | Tapp | 1.8 | 2.5 | 3.6 |
|  |  |  | No Tapp | 1.1 | 1.8 | 3.2 |
|  |  | Distance | Tapp | 2.9 | 3.3 | 4.3 |
|  |  |  | No Tapp | 3.8 | 4.9 | 7.5 |
| Nicotine | Nicotinic acetylcholine receptor agonist | Thigmotaxis | Tapp | 4.1 | 6.2 | 6.3 |
|  |  |  | No Tapp | Model unusable |  |  |
|  |  | Distance | Tapp | 0.5 | 0.70 | 0.9 |
|  |  |  | No Tapp | Model unusable |  |  |
| Dexamethasone | Glucocorticoid receptor agonist | Thigmotaxis | Tapp | 0.2 | 0.31 | 0.9 |
|  |  |  | No Tapp | 0.08 | 0.3 | 0.8 |
|  |  | Distance | Tapp | Non-significant. Analysis not done |  |  |
|  |  |  | No Tapp | Non-significant. Analysis not done |  |  |
| Ethylenethiourea | Thyroid-disrupting goitrogen | Thigmotaxis | Tapp | 8.3 | 22.0 | 22.9 |
|  |  |  | No Tapp | 15.5 | 38.1 | NC |
|  |  | Distance | Tapp | Model unusable |  |  |
|  |  |  | No Tapp | Model unusable |  |  |

For visual stimulus experiments, exposure to Chlorpyrifos caused an increase in thigmotaxis during dark and light periods that was significant for all studied concentrations (Fig. 5A) with a BMD between 0.3-0.4 mg/L (Table 3). On the other hand, total distance moved was also affected by an increase in activity during both visual periods but exacerbated during light intervals (Fig. 5B). A significant response was not observed at lower concentrations and BMDs were 1-2 orders of magnitude higher (3-43 mg/L; Table 3). Moreover, at highest concentrations, the trend was reversed with a decrease in distance moved to control levels even though thigmotaxis continued to be increased (Fig. 5 A vs B). Similarly, larvae treated with Nicotine exhibited a significant increase in thigmotaxis that started at the lowest concentration tested for dark periods (Fig. 5C) with a BMD of 0.15 mg/L. Total distance moved during dark periods also exhibited a significant increase but only at the highest concentrations tested with a 5 times higher BMD of 0.8 mg/L, and this trend was reversed at highest concentrations Figs 5D). Thigmotaxis response to Nicotine during light periods was less pronounced and only significant for the two highest concentrations (Fig. 5C). No significant differences were observed for distance moved during light periods (Fig. 5D). Exposure to all concentrations of Dexamethasone induced a significant thigmotaxis increase in larvae during both visual stimulus periods although more pronounced, with lower BMD, for light intervals (Fig. 5E, Table 3). On the contrary, no significant changes in activity were observed (Fig. 5F). Ethylenethiourea also induced thigmotaxis in exposed larvae but in this case the effect was more pronounced during dark intervals where significant changes were observed even at the lowest concentrations (Fig. 5G). Contrastingly, distance moved was not significantly altered during dark periods and only slightly and non-monotonically altered during light intervals (Fig. 5H). Moreover, BMD for distance moved was two orders of magnitude greater than the one for thigmotaxis (Table 3).

Response to the negative non-neuroactive substances Saccharin and Amoxicillin was as expected, with no significant changes in thigmotaxis or distance travelled (Fig. 6), except for a significant increase in both parameters for Amoxicillin at the highest concentration (Fig. 6C, D).

**Fig 6.**
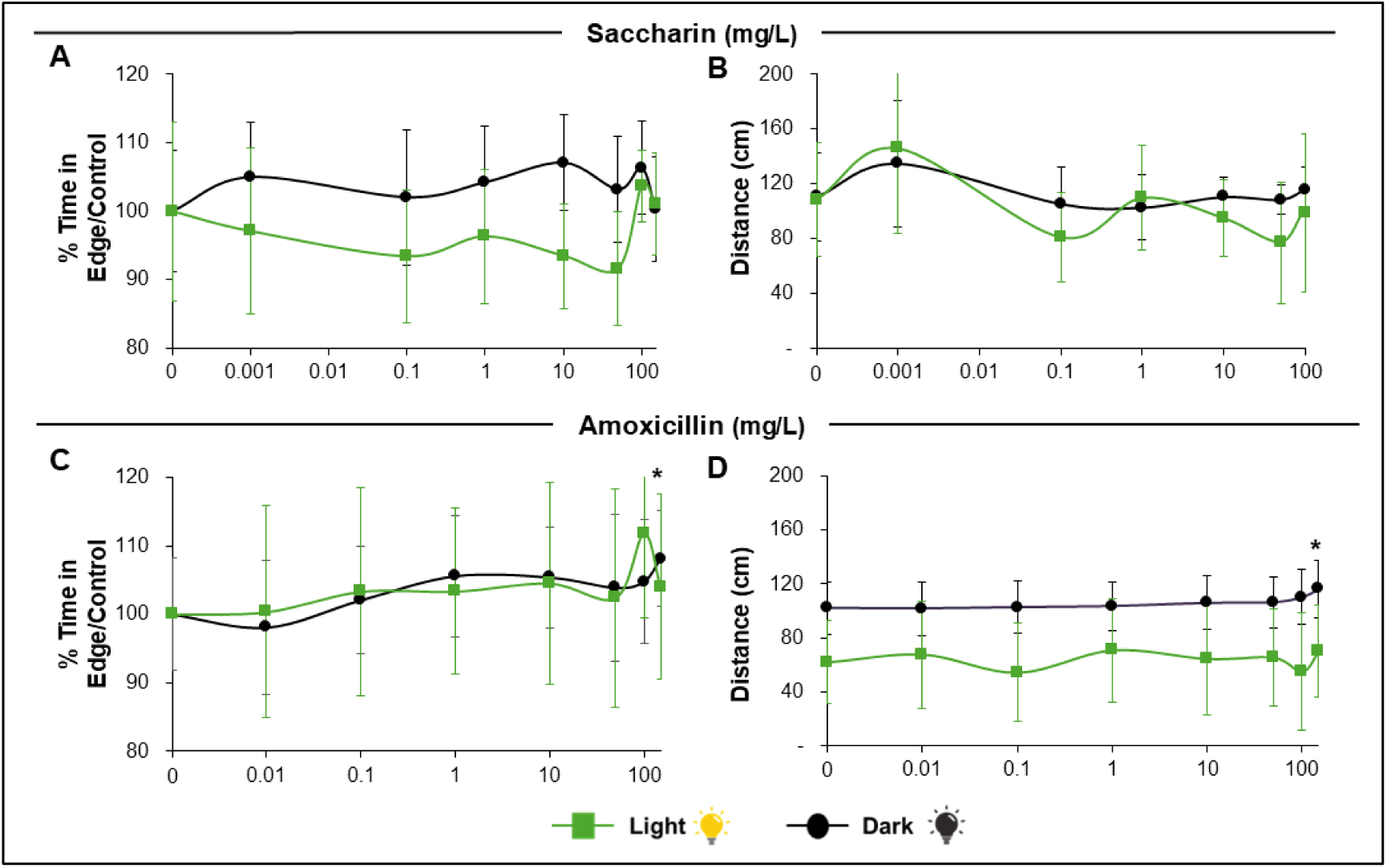
Results of Thigmotaxis and Total distance moved by larvae exposed to negative non-neuroactive substances (saccharin and amoxicillin) in 96S well plates after visual stimulation. Green lines depict light periods, and black lines depict dark periods. Green asterisk above point denotes significant differences of substance concentration vs. control during light period. Black asterisk above point denotes significant difference of substance concentration vs. control during dark period. Data are presented as Means +/- SD.

### 3.4. Neuroactive and negative substances acoustic tests

Acoustic stimulation also had dissimilar effects on thigmotaxis and distance moved that depended on the substance studied. Chlorpyrifos exposure caused an increase in % time in edge during tapping and quiet (no tapp) periods that was significant starting at 3 mg/L (Fig. 7A) with a BMD between 1.8-2.5 mg/L (Table 3). Similarly, total distance moved was affected by an increase in activity during both tapping and quiet periods although a significant response was only observed starting at 10 mg/L (Fig. 5B) and BMDs were higher, especially for response during quiet periods (1.8 mg/L for thigmotaxis vs 4.9 mg/L for distance; Table 3). Similarly to results during visual stimulation, the largest concentrations exerted an inverse effect with mean distance moved returning to control levels whereas thigmotaxis continued to be altered (Fig. 7 A vs. B).

**Fig 7.**
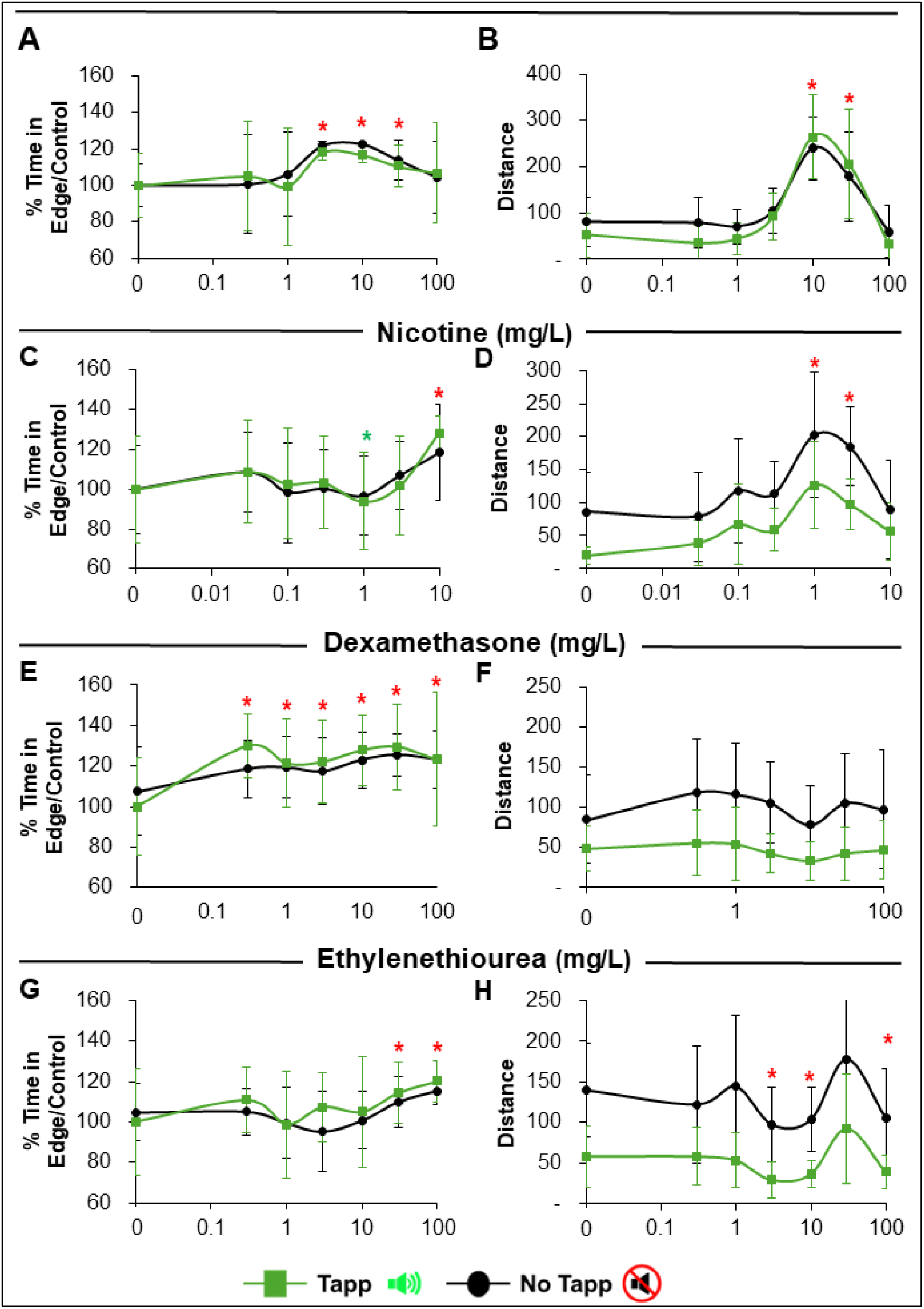
Results of Thigmotaxis and Total distance moved by larvae exposed to neurotoxic substances (chlorpyriphos, nicotine, dexamethasone and ethylenethiourea) in 96S well plates after acoustic stimulation. Green lines depict tapping periods and black lines depict quite (no tapping) periods. Green asterisks above point denote significant differences of substance concentration vs control during tapping period. Black asterisk above points denotes significant difference of substance concentration vs control during quiet period. Red asterisks above points denote significant differences of substance concentration vs. control for both periods. Data are presented as Means +/- SD.

Larvae treated with Nicotine exhibited a non-monotonic response with a significant decrease of thigmotaxis at 1 mg/L and an increase at 10 mg/L, for both tapping and silent periods (Fig. 7C). On the other hand, even though there is a trend of increase in distance moved, this was not statistically significant during tapping periods and only significant at 1 and 3 mg/L for quiet periods (Fig. 7D). This trend was revered at highest concentration tested (Fig. 7D) for distance moved but not for thigmotaxis. Exposure to all concentrations of Dexamethasone induced a significant thigmotaxis increase in larvae during both acoustic stimulus periods with similar BMDs of ∼ 0.3 mg/L for both intervals (Fig. 7E, Table 3). Contrarily, no significant changes in activity were observed (Fig. 7F). Ethylenethiourea induced thigmotaxis in exposed larvae but only significantly at the two highest concentrations with BMDs of 22 and 38 mg/L for quiet and tapping periods respectively (Fig. 7G, Table 3). On the other hand, distance moved exhibited a non-monotonic and highly variable response with a significant decrease at concentrations of 3, 10, and 100 mg/L and an increase at 30 mg/L during both quiet and tapping periods (Fig. 7H). Regarding the negative substances Saccharin and Amoxicillin, results showed no significant changes in thigmotaxis or distance travelled during both tapping and quiet periods for all studied concentrations (up to 100 mg/L; Fig. 8).

**Fig 8.**
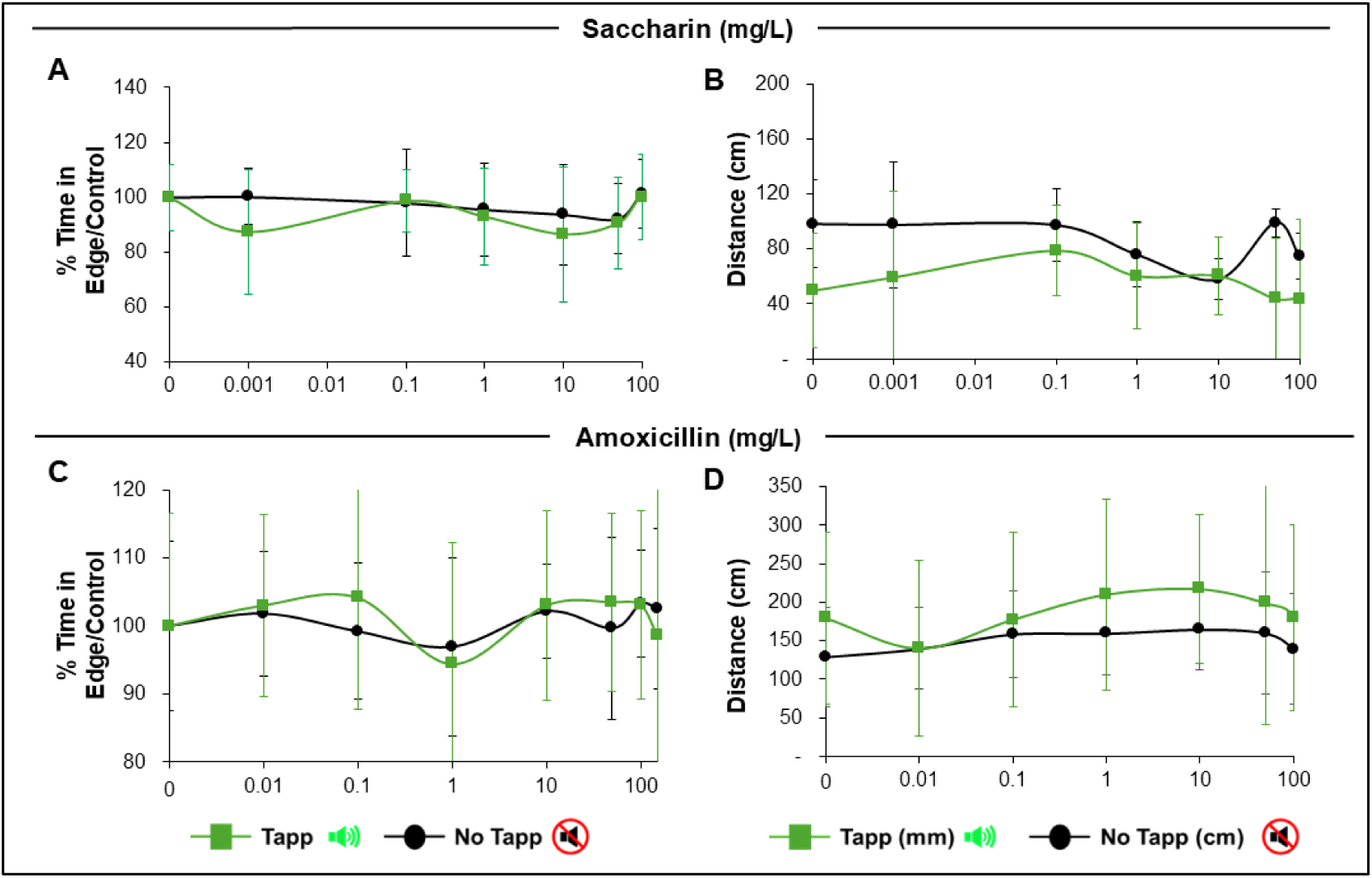
Results of Thigmotaxis and Total distance moved by larvae exposed to negative non-neuroactive substances (saccharin and amoxicillin) in 96S well plates after acoustic stimulation. Green lines depict tapping periods and black lines depict quite (no tapping) periods. No significant differences were found. Data are presented as Means +/- SD.

## 4. Discussion

Behavioral studies remain crucial in evaluating developmental neurotoxicity due to difficulties in directly detecting effects on particularly affected nervous system regions (Heyer and Meredith, 2017; Vorhees et al., 2021). In this context, the goal of the present study was the development of a high throughput thigmotaxis assay to assess anxiety-like behavior in zebrafish larvae at 120 hpf suitable for studying effects of toxic substances and pharmaceuticals for both the ecotoxicology and human toxicology fields. Our results provide evidence that this assay generates results comparable to the ones obtained with the traditional lower throughput method. Moreover, we have shown that the high throughput assay is sensitive to anxiogenic and anxiolytic compounds and, by incorporating both visual and acoustic stimulation, it offers a comprehensive method for evaluating different neuroactive substances under varied conditions.

During assay development, our first objective was to evaluate the performance of the 96S well plate compared to the 24 round well plate format commonly used (Merola et al., 2021a; Schnörr et al., 2012) in order to increase throughput. Our results show that both configurations gave very similar results both in control larvae behavior and also when larvae were exposed acutely (1h) to the model anxiogenic Caffeine and the model anxiolytic Diazepam, regardless of the type of stimulus present (Figs 3, 4). These similarities were particularly important for thigmotaxis (% time at the edge of well) even though minor differences in locomotion (distance moved) were observed during light periods, probably due to the greater surface area of 24R wells. To confirm these results, we performed a TOST analysis (Fig. A.2 and Table A.1). We chose a conservative prespecified equivalence margin of Δ = ±10 pp, which was justified by the variability of the assay (SD approx. 10 – 30 pp) and the biological interpretability of the thigmotaxis assay readings. The analysis showed practical equivalence for the majority of assay and condition contrasts, including controls for Visual Time% and Distance cm and Acoustic Time% and Distance cm, all caffeine conditions, and diazepam under Visual Dark and Acoustic Tap. Only a few comparisons were not equivalent which likely reflects endpoint-specific scaling rather than systematic format bias. Considering all of the above and given that the 96S well plate format supports higher throughput and automatization and is compatible with image analysis, we strongly support the use of this format for evaluating anxiety-like behavior in zebrafish larvae as it is suitable for regulatory applications without compromising data integrity (Gerlai, 2010).

A second step was to test the sensitivity of the 96S plate to thigmotaxis changes in response to known anxiotropic substances. Our results show that larvae in this plate format responded in general as expected considering the pharmacological profiles of Caffeine and Diazepam (Fig. 4). Response to Caffeine, a known adenosine receptor antagonist (Ribeiro and Sebastião, 2010), was a robust increase in anxiety-like behavior with greater thigmotaxis during light and acoustic stimulus periods but not during dark periods. The documentation of stimuli-dependent responses is important as few works have studied these sensory modulated responses in zebrafish larvae, most reporting caffeine-related thigmotaxis data only during dark or light periods without transitions (Richendrfer et al., 2012a; Schnörr et al., 2012; Tan et al., 2022). On the other hand, the anxiolytic response to the GABAA receptor agonist Diazepam was only observed during dark periods in accordance with previous research done with larval zebrafish (Widelski et al., 2021). During light and acoustic periods there was a lack of response at lower concentrations and an increase in anxiety at the two highest concentrations. Even though it has been documented that the GABA switch from excitatory to inhibitory can occur in certain regions of the zebrafish brain as early as 2.120 hpf (Zhang et al., 2010), our results suggest that the switch might not be synchronous and that some brain areas, related to acoustic responses for example, might still have excitatory GABAergic neurons (Kirmse et al., 2018). This is further supported by the increased anxiety observed (Fig. A.2) when 120 hpf larvae were exposed to Valerenic acid, a known GABA receptor agonist (Benke et al., 2009). In addition, other behavioral modulating pathways could be playing a role as benzodiazepines have been reported to also affect the glutamatergic, dopaminergic, serotonergic and noradrenergic pathways (Gomez-A et al., 2017; Kellogg and Retell, 1986; Lista et al., 1989; Zeller et al., 2008). These results suggest a complex modulation of the anxiety response depending on the processing of sensory inputs and highlights the importance of conducting thigmotaxis assays that challenge the larvae with multiple stimuli in order to better detect possible neuroactive substances and help elucidate mechanisms of action.

Mean total distance moved in light and/or dark periods is the most common method of assessing behavioral alterations in zebrafish larvae exposed to neurotoxicants and pharmacological compounds (Dach et al., 2018; Haigis et al., 2022; Irons et al., 2010). In this study, we compared results obtained by this method to results obtained by our thigmotaxis method in larvae that were exposed to known neurotoxicants and negative substances as a proof-of-concept approach. In general, thigmotaxis proved to be a more sensitive and discriminating endpoint than mean distance moved.

Exposure to Chlorpyrifos, a known acetylcholinesterase inhibitor (Yen et al., 2011), had a clear effect on zebrafish larvae behavior, as had been shown before (Levin et al., 2004; Silva, 2020). Moreover, our results of mean distance moved by exposed larvae during light and dark periods are similar to those obtained previously (Quevedo et al., 2018), showing overall hyperactivity (during visual and acoustic stimuli) and loss of the normal pattern of less movement in the light and more in the dark (Figs. 5, 6B). However, significant effects were observed starting at higher doses than those observed by the thigmotaxis endpoint (Figs. 5, 6A), and benchmark doses were generally lower for the edge preference endpoint. A similar response was observed for the acoustic stimulus test, with thigmotaxis being more sensitive, particularly during the quiet periods. All this suggests that cholinergic hyperstimulation provoked at early stages may induce anxiety-like behavior without altering general locomotion at low doses.

Similarly, exposure to Nicotine, a nicotinic acetylcholine receptor (nAChR) agonist (Tiwari et al., 2020), elicited an increase in larval activity at lower concentrations, especially during dark (Fig. 5D) and silent (Fig. 7D) periods. This was followed by a sharp decrease to control levels at higher concentrations, as expected due to previous literature (Petzold et al., 2009). On the other hand, Nicotine elicited increased thigmotaxis, especially during dark periods (Fig. 5C) or had a U-shaped response with a decrease edge preference at low concentrations during acoustic stimulation (Fig. 7C). As in mammalian models (Cohen et al., 2009), adult zebrafish responses to acute Nicotine exposure have been found to be mainly anxiolytic although anxiogenic behaviors have also been observed, especially for chronic exposures (Wronikowska et al., 2020) whereas acutely exposed larvae studies are scarce but have also observed increased thigmotaxis with similar doses to the present study (Chen and Scalzo, 2015). In zebrafish, as in mammals (Ackerman et al., 2009), nicotine can exert its mode of action through various neuroregulatory pathways (GABA, dopamine, norepinephrine, serotonin) as nicotinic acetylcholine receptors can be found in different areas of the nervous system (Tiwari et al., 2020). Evidently, complex neurotransmitter pathways are playing a role, but it was not the subject of this work to disentangle molecular mechanisms of action. Nevertheless, our results show that nicotine is affecting larva zebrafish differently than typical responses in adults and behavior varied depending on type of stimulus as anxiolytic effects were indeed observed during the acoustic test. Most importantly, thigmotaxis was again a more sensitive endpoint to evaluate neuroactivity than mean distance moved, especially for the light/dark transition test.

Dexamethasone is a synthetic glucocorticoid receptor agonist used in humans as an anti-inflammatory that has been shown to increase anxiety behaviors in rodents (Onaolapo et al., 2014) and zebrafish (Khor et al., 2013). Therefore, it was not surprising to observe increased edge preference in treated larvae although the degree of effect varied with stimuli type with the most thigmotaxis present during light periods (Fig. 5E). In contrast, distance moved was largely unaffected (Figs 5, 7F). Therefore, our results suggest that dexamethasone is activating the hypothalamic-pituitary-adrenal axis affecting stress-related pathways without affecting general locomotion-related circuits (McLean et al., 2008). This is interesting as previous work has documented altered locomotion in addition to anxiety in 72-120 hpf exposed zebrafish larvae (Khor et al., 2013). It is possible that this difference is due to different exposure regimes as concentration ranges were similar. Nevertheless, in this case the thigmotaxis assay was again more sensitive in detecting neuroactivity under both visual and acoustic stimulation.

Ethylenethiourea is an organosulfur compound formed by degradation of dithiocarbamate fungicides (Stadler et al., 2022). It is a known thyroid disruptor (Ma et al., 2024) decreasing thyroid hormones in exposed mammals (Nebbia and Fink-Gremmels, 1996) and zebrafish (Thienpont et al., 2011) by inhibiting the enzyme thyroid peroxidase (Doerge and Takazawa, 1990). Moreover, it has been shown to exhibit neurotoxic effects (Bjørling-Poulsen et al., 2008) by directly promoting neuronal degeneration (Khera, 1987). Similarly, neurotoxicity has been observed to be elicited by other dithiocarbamate compounds by altering glutamate transport (Vaccari et al., 1999) or affecting mitochondria (Domico et al., 2006). Our results indicate that ethylenethiourea altered both thigmotaxis and general locomotion responses although effects varied depending on type of stimulus. During dark intervals ethylenethiourea exposure increased thigmotactic behavior in larvae even at the lowest concentration tested although time in edge also increased to a lesser degree in light intervals. In this case larvae also exhibited hyperlocomotion but only during dark periods (Fig. 5G, H). On the other hand, sound stimulation elicited thigmotaxis only at the highest concentrations tested and activity showed a non-monotonic response with hypoactivity detected at intermediate concentrations (Fig. 7G, H). Increase in anxiety-related responses as a result of ethylenethiourea exposure could be explained by previously observed effects on glutamate transport as extracellular glutamate accumulation could lead to neuronal hyperexcitability (McKeown et al., 2012). This could also explain the observed hyperlocomotion in dark periods but not the hypolocomotion observed during quite periods after sound stimulation although it is possible that neurotransmitter pathway alterations affect visual and acoustic circuits differently (Pfeffer et al., 2021). In the case of ethylenethiourea, again thigmotaxis proved to be a more sensitive endpoint for detecting behavioral alterations, especially during periods of visual stimulation.

In contrast, exposure to the non-caloric sweetener Saccharin did not affect significantly the thigmotaxis or general locomotion in exposed larvae, validating its role as a negative control. The beta-lactam antibiotic Amoxicillin could also serve as a negative control as it only slightly altered locomotion and edge preference at the highest concentration tested. Since beta-lactam antibiotics have been observed to modulate glutamatergic signaling (Rothstein et al., 2005), this effect could reflect these drugs’ unintended neuroactive properties at high doses. However, these responses were not consistent across stimuli suggesting a weak or secondary effect.

In summary, our findings have proven that increasing the thigmotaxis assay throughput from 24R to 96S well plates does not affect performance, making it a great candidate for drug and toxicant screening of neuroactive substances. Our work provides further evidence of thigmotaxis being pharmacologically validated as an anxiety-related behavior in zebrafish larvae (Schnörr et al., 2012; Richendrfer et al., 2012). Nevertheless, while these behavioral responses support an anxiety-like interpretation, direct coupling with molecular endpoints such as cortisol or neurotransmitter levels has not been consistently established in 120 hpf larvae (Best and Vijayan, 2018). Nevertheless, edge-preference behavior is a window into the complex functioning of the nervous system involving sensory processing, motor coordination and decision making. This work has demonstrated that thigmotaxis is a valuable and sensitive endpoint in studying effects of neuroactive substances. Moreover, our results highlight the greater sensitivity of this edge-preference assay that includes both acoustic and visual stimuli compared to the more commonly used general locomotion Light/Dark transition test. However, we believe that the integration of both endpoints could be valuable, especially when integrating stimulus-specific thigmotaxis with locomotion to refine the detection of compound-specific behavioral signatures that could inform hypothesis of possible modes of action (Fig. 9). Notably, substances like ETU and Nicotine showed differential responses under visual and acoustic stimulation, suggesting different neural pathways are involved. Nonetheless, while thigmotaxis has shown to be more sensitive than total distance moved in most cases, the interpretation of results can be complex due to overlapping neurotransmitter systems and potential developmental variability in receptor expression. Moreover, caution should be taken as larval developmental stage could also play a role and results obtained at 120 hpf might not be directly extrapolatable to other, more advanced larval stages or adult individuals. Nevertheless, although developmental differences may influence edge-preference outcomes, the 120 hpf thigmotaxis assay remains a sensitive indicator of neuroactivity and a promising approach to elucidate potential modes of action when integrated with molecular analyses.

**Fig 9.**
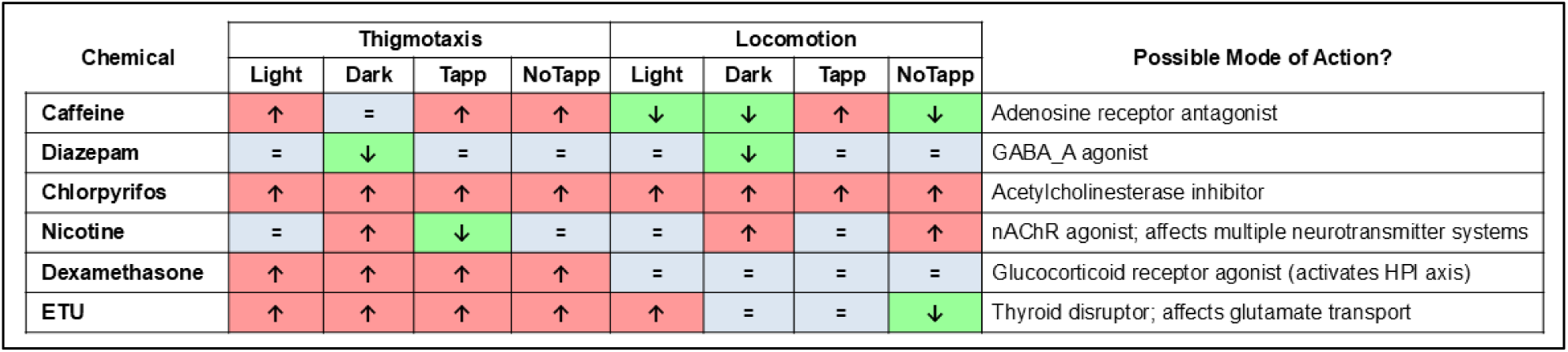
Summary of effects caused by model substances to larvae exposed in 96S well plates in visual (Light/Dark) or acoustic (Tapp/No Tapp) modes. Th: Thigmotaxis (% time in edge); Lo: Locomotion (total distance moved). Data on distance moved for caffeine and diazepam are taken from Fig. A.4

In light of our results, we propose the designation Z-MULTITHIGMO NAM (Zebrafish Multiplexed Thigmotaxis New Approach Methodology) to describe our multiplexed larval workflow integrating visual and acoustic stimuli within a standardized thigmotaxis paradigm (Fig. 10). The assay can be positioned either as a stand-alone new approach methodology or as a plug-in module within existing behavioral NAM batteries (e.g., photomotor and visual–acoustic paradigms). Its main strengths include: (i) construct validity demonstrated with the reference pair—anxiogenic caffeine and anxiolytic diazepam; (ii) mechanistic coverage across major neurochemical axes, as shown with probes targeting cholinesterase inhibition (chlorpyrifos), nicotinic receptor activation (nicotine), glucocorticoid signaling (dexamethasone), and a thyroid-disrupting negative control (ethylenethiourea), together with putative negatives (saccharin, amoxicillin); (iii) robustness across sensory modalities (visual and acoustic) and assay formats (24- and 96-well), demonstrating practical equivalence between both configurations; and (iv) quantitative potency metrics (BMD/BMDL/BMDU) consistently showing higher sensitivity for thigmotaxis than for locomotor activity, thereby enabling potency ranking and mechanistic read-across.

**Fig 10.**
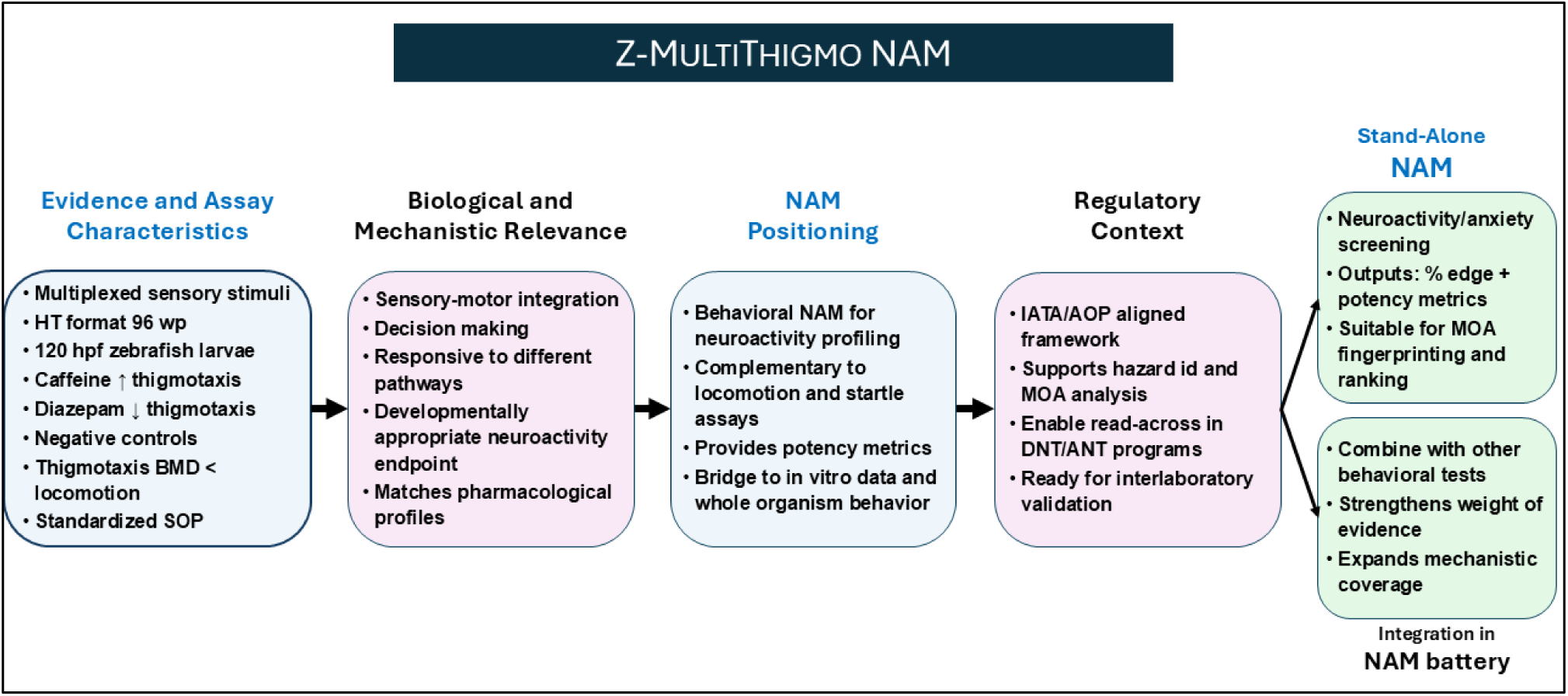
Positioning of the Z-MULTITHIGMO assay within the NAM landscape. Z-MULTITHIGMO is a multiplexed behavioral thigmotaxis assay in zebrafish larvae using visual and acoustic stimuli. Implemented in 96-well high-throughput (HT) format under a versioned standard operating procedure (SOP), it shows predicted pharmacology (caffeine anxiogenic, diazepam anxiolytic) and delivers quantitative benchmark dose (BMD) metrics for potency ranking and mode of action (MOA) interpretation. Within the New Approach Methodology (NAM) framework, the assay aligns with Integrated Approaches to Testing and Assessment (IATA) and Adverse Outcome Pathway (AOP) concepts, and complements photomotor, light/dark, and startle/habituation tests. Its intended context of use spans developmental neurotoxicity (DNT) and adult neurotoxicity (ANT) screening, with readiness for inter-laboratory validation.

Compared with other behavioral NAMs, the Z-MULTITHIGMO NAM adds value by combining stimulus multiplexing, standardized arena geometry, and explicit BMD modeling within a single harmonized standard operating procedure (SOP). This integration improves coherence and reproducibility and aligns the assay with current EU strategies to advance DNT/ANT assessment within Integrated Approaches to Testing and Assessment (IATA) and Adverse Outcome Pathway (AOP) contexts. Forthcoming steps should focus on conducting multi-laboratory ring trials to establish inter-laboratory reproducibility and delineate the assay’s domain of applicability.

When framed within its defined context of use (screening-level neurotoxicity or neuromodulation hazard identification with potency ranking) the Z-MULTITHIGMO NAM is ready to function as a behavior-based NAM. In addition, it has entered a collaborative evaluation phase within the PARC neurotoxicity group, where a blind set of reference chemicals is being tested across partner laboratories. This exercise will define its role within a future adult neurotoxicity (ANT) in vitro battery. In parallel, a developmental neurotoxicity (DNT) version of the assay is being established, extending exposure from 2 hpf onward to capture early neurodevelopmental perturbations and enable direct comparison with the current ANT mode.

In summary, this study demonstrates that this multi-stimuli zebrafish larvae thigmotaxis assay is a robust, reproducible, and sensitive method for the detection of neuroactive substances. We have demonstrated that the assay is scalable to a high throughput format (96 well plates) and that adding visual and acoustic stimulation is important to increase sensitivity and to distinguish compounds’ MOA, making it more sensible than general locomotion measures. Altogether the Z-MULTITHIGMO NAM is a practical and ethical assay that is mechanistically informative, making it a valuable tool to contribute to neurotoxicity evaluation that would be valuable for ecotoxicology as well as to assess effects on the human nervous system.

## CRediT authorship contribution statement

**Torres-Ruiz M.:** Resources, Methodology, Investigation, Funding acquisition, Formal analysis, Data analysis, Data curation, Writing – review & editing, Writing – original draft. **Muñoz Palencia M**.: Methodology, Investigation, Data analysis, Data curation. **De la Vieja A.:** Resources, Funding acquisition, Formal analysis, Data analysis, Data curation, Figure editing, Writing – review & editing. **Cañas-Portilla A.:** Resources, Funding acquisition, Writing – review & editing.

## Compliance with ethical requirements

This article does not contain any studies on human or animal subjects.

## Acknowledgements

We thank all members of PARC WP 5.2.1e for their suggestions and insights regarding the development of this assay.

## Data availability

Data will be made available on request

## Appendix A. Supplementary Data

Figure A.1: Tests of solvent (DMSO) concentrations to evaluate maximum doses in thigmotaxis assay in both visual and acoustic. Figure A.2. Representation of 96S–24R plate differences using Forest/TOST plots. Figure A.3: Plate-format comparison for % time at edge relative to control in 120 hpf larvae for model substances. Figure A.4: Results of total distance moved by larvae exposed to Caffeine and Diazepam in 96S well plates after visual (left) and acoustic (right) stimulation

## SUPPLEMENTARY MATERIAL

**Figure A.1:**
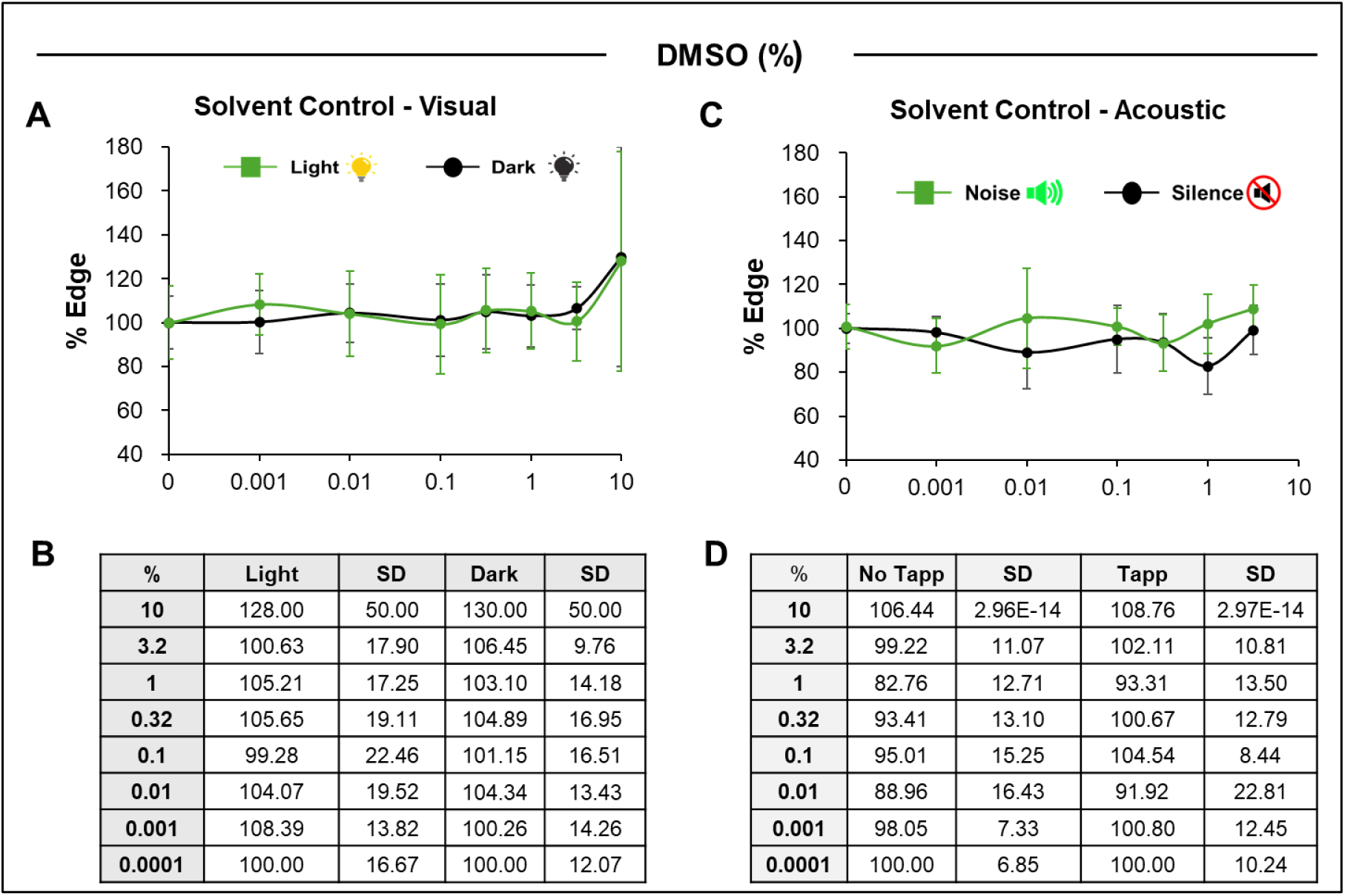
Tests of solvent (DMSO) concentrations to evaluate maximum doses in thigmotaxis assay in both visual and acoustic (n=12) modes in 24/96 well plates. Panel (A) and (B) depict graphic representation and values of the percent distance in the edge by the larvae for the edge of the plates after visual stimuli (light in green, n=36; dark in black, n=36). Panel (C) and (D) depict graphic representation and values of the percent distance in the edge by the larvae for the edge of the plates after acoustic (noise in green, n=12; silence in black, n=12).

**Figure A.2.**
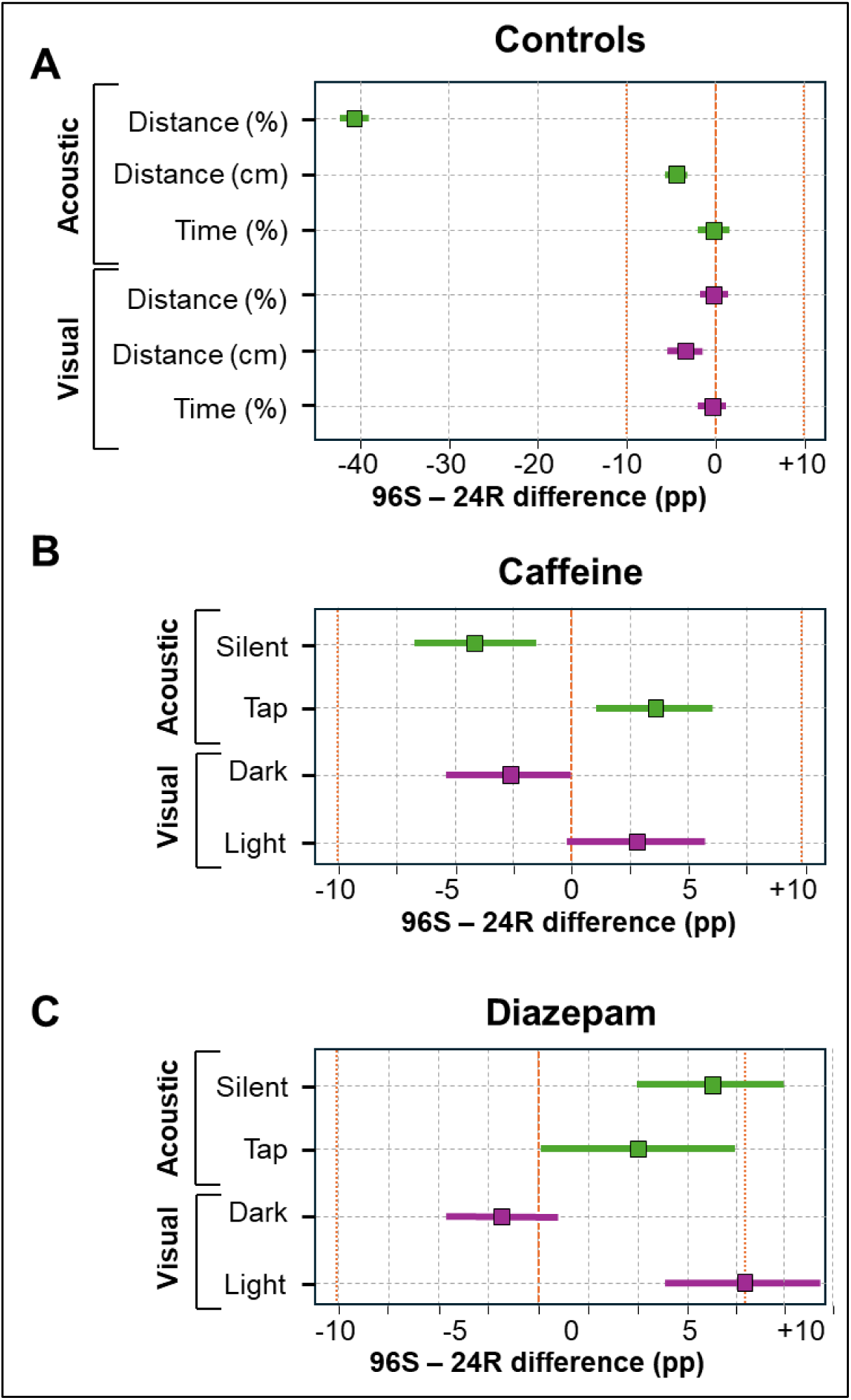
Representation of 96S–24R plate differences using Forest/TOST plots. (Δ = ±10 percentage points). Each point shows the pooled mean difference (96S − 24R) and the horizontal bar is the 90% CI. The vertical orange dashed line marks 0 (no difference) and the orange dotted lines mark the equivalence bounds at ±Δ (±10 percentage points).. A) Controls endpoints include Time (%), Distance (%), and Distance (cm) at Edge. Visual stimuli are Light/Dark and acoustic stimuli are Tap/Silent (pooled). Caffeine (B) and diazepam (C) endpoints are Time % (Edge). Values are pooled across concentrations within each assay/condition using fixed-effect inverse-variance meta-analysis with Welch/Satterthwaite uncertainty. A comparison is classified as Equivalent when the entire 90% CI lies within ±Δ; if the CI extends beyond these bounds, the result is not equivalent and is further labeled Different when the two-sided Welch test is significant (p < 0.05), otherwise Inconclusive. Abbreviations: TOST, two one-sided tests; pp, percentage points; CI, confidence interval; Welch, unequal-variance t-test.

**Figure A.3:**
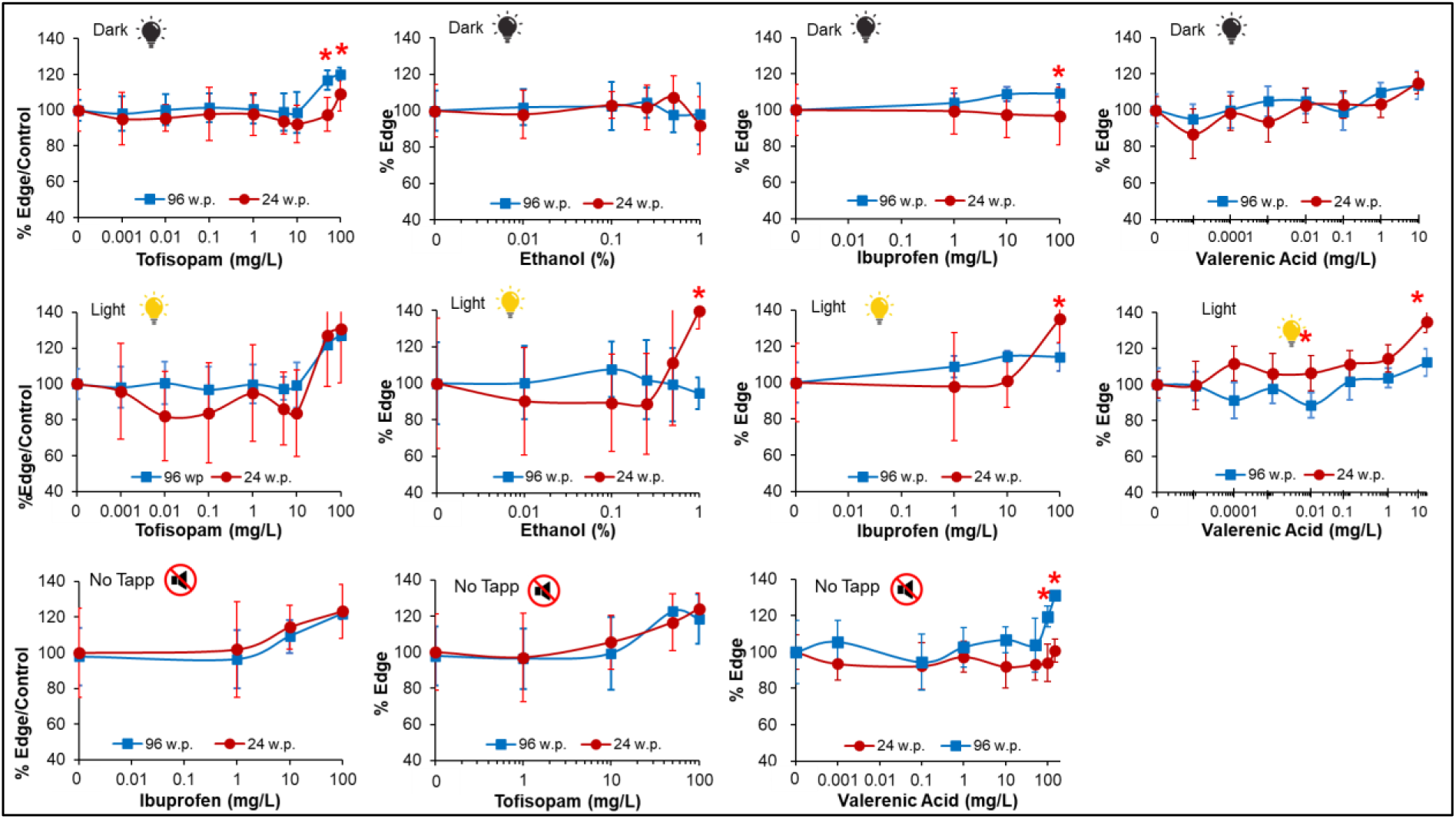
Plate-format comparison for % time at edge (thigmotaxis) relative to control in 120 hpf larvae. Larvae were exposed for 1 h at 120 hpf and tested in 96-well square plates (96S; blue) and 24-well round plates (24R; red). Curves show the percentage of time spent at the edge normalized to the corresponding control for each concentration. For tofisopam, ibuprofen, and valerenic acid, responses are shown during dark (dark light-bulb icon), light (light-bulb icon), and silence following tapping (muted-speaker icon) periods. For ethanol, only dark and light periods are displayed. Symbols indicate mean ± SD. Asterisks above points denote significant differences between plate formats at the same concentration and stimulus (p ≤ 0.05).

**Figure A.4:**
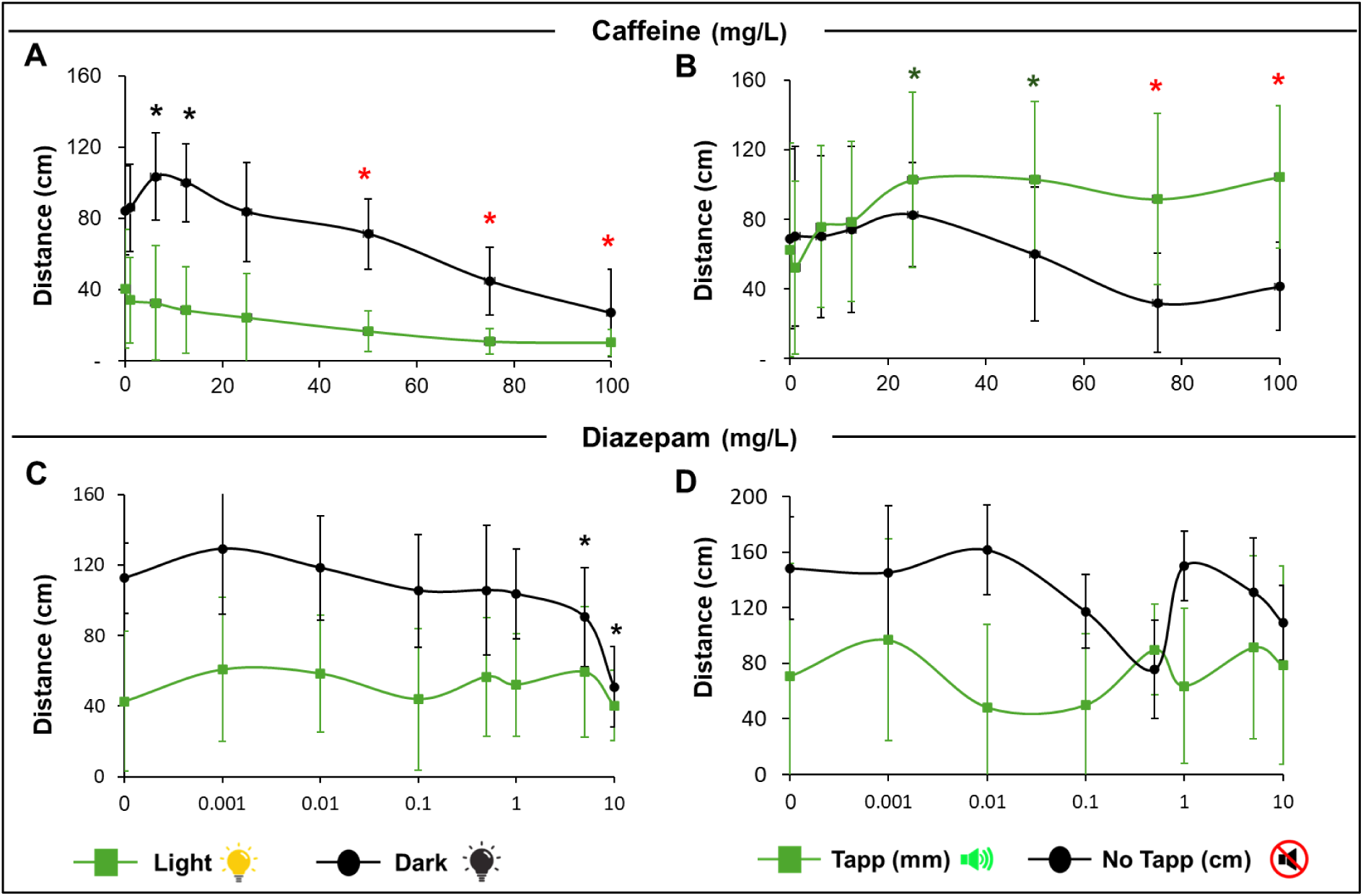
Results of total distance moved by larvae exposed to Caffeine (top panels) and Diazepam (bottom panels) in 96S well plates after visual (left) and acoustic (right) stimulation. Green lines depict light or tapping periods and black lines depict dark or quite (No tapping) periods. Green asterisks above point denote significant differences of substance concentration vs control during light/tapping period. Black asterisk above points denotes significant difference of substance concentration vs control during dark/quiet period. Red asterisks above points denote significant differences of substance concentration vs. control for both periods. Data are presented as Means +/- SD.

